# Dilated cardiomyopathy-associated RNA Binding Motif Protein 20 regulates long pre-mRNAs in neurons

**DOI:** 10.1101/2023.12.06.570345

**Authors:** Giulia Di Bartolomei, Raul Ortiz, Dietmar Schreiner, Susanne Falkner, Esther E. Creemers, Peter Scheiffele

## Abstract

Precise coordination of molecular programs and neuronal growth govern the formation, maintenance, and adaptation of neuronal circuits. RNA metabolism has emerged as a key regulatory node of neural development and nervous system pathologies. To uncover cell-type-specific RNA regulators, we systematically investigated expression of RNA recognition motif-containing proteins in the mouse neocortex. Surprisingly, we found RBM20, an alternative splicing regulator associated with dilated cardiomyopathy, to be expressed in cortical parvalbumin interneurons and mitral cells of the olfactory bulb. Genome-wide mapping of RBM20 target mRNAs revealed that neuronal RBM20 binds pre-mRNAs in distal intronic regions. Loss of neuronal RBM20 has only modest impact on alternative splice isoforms but results in a significant reduction in an array of mature mRNAs in the neuronal cytoplasm. This phenotype is particularly pronounced for genes with long introns that encode synaptic proteins. We hypothesize that RBM20 ensures fidelity of pre-mRNA splicing by suppressing non-productive splicing events in long neuronal genes. This work highlights a common requirement for RBM20-dependent transcriptome regulation in cardiomyocytes and neurons and demonstrates that a major genetic risk factor of heart disease impacts neuronal gene expression.

## Introduction

Neurons exhibit complex transcriptional programs that instruct the specification of functionally distinct neuronal cell types. Functional and anatomical properties of neurons emerge during development. However, the molecular underpinnings that define cell type-specific properties are only beginning to be elucidated. RNA-binding proteins (RBPs) have emerged as key regulators of neuronal function through modification of mRNA processing, localization, stability, and translation (Ule & Darnell, 2006; Babitzke *et al*, 2009; Vuong *et al*, 2016; Mauger & Scheiffele, 2017; Holt *et al*, 2019; Ule & Blencowe, 2019; Gomez *et al*, 2021). Moreover, RBP dysfunction is a significant contributor to pathologies, including neurodevelopmental and neurodegenerative conditions (Ling *et al*, 2013; Lopez Soto *et al*, 2019; Gebauer *et al*, 2021; Schieweck *et al*, 2021). Here, we discovered an unexpected neuronal function for the SR-related protein, RNA Binding Motif Protein 20 (RBM20). Thus far, RBM20 was considered to be muscle-specific and to represent a key alternative splicing regulator in cardiomyocytes. Mutations in the *RBM20* gene are linked to an aggressive form of dilated cardiomyopathy (Brauch *et al*, 2009; Parikh *et al*, 2019). In cardiomyocytes, RBM20 protein controls alternative exon usage of transcripts encoding key sarcomere components (Titin and Tropomyosin) and proteins involved in calcium signaling, such as CAMK2D and the α1 subunit of the L-type voltage gated calcium channel (CACNA1C) (Guo *et al*, 2012; Maatz *et al*, 2014; van den Hoogenhof *et al*, 2018; Zhu *et al*, 2021). RBM20 contains an RNA Recognition Motif (RRM) domain, two zinc finger motifs, and an extended arginine/serine-rich region. This domain organization is shared with the paralogues Matrin-3 (MATR3) and ZNF638 (Watanabe *et al*, 2018), and is similar to FUS and TDP43, two RNA/DNA binding proteins that regulate various steps of RNA metabolism. Comprehensive RBM20 protein-RNA interaction maps have been defined for cardiomyocytes (Maatz *et al*., 2014; van den Hoogenhof *et al*., 2018; Briganti *et al*, 2020). However, RBM20 expression and function in the brain, are unknown. Considering its highly selective expression in specific cell populations and the critical roles in cardiomyocytes, we hypothesized that neuronal RBM20 controls key steps of RNA metabolism and contributes to the regulation of neuronal gene expression.

## Results

### RBM20 is selectively expressed in specific GABAergic and glutamatergic neurons

To discover regulators of neuronal cell type-specific transcriptomes, we analyzed a neuronal gene expression dataset covering cortical pyramidal cells and the major GABAergic interneuron populations of the mouse cortex (Furlanis *et al*, 2019). We generated a hand-curated list of 234 potential RBPs, selected based on the presence of at least one predicted RRM domain (Fig. 1A and Fig. S1A). Transcripts for 182 of these RRM proteins were significantly expressed in mouse cortical neurons. While some (such as *Eif3g*, *Hnrnpl*) were broadly detected across all neuronal populations, others were differentially expressed in specific neuron classes. Thus, *Rbm38* was almost exclusively expressed in VIP^+^ (vasoactive intestinal polypeptide-positive) interneurons and *Ppargc1a and Rbm20* in parvalbumin-positive (PV^+^) interneurons, indicating that they may play an important role in cell type-specific gene regulation. Identification of *Rbm20* expression in the brain was surprising considering that the RBM20 protein was thought to be muscle-specific. Thus, we focused our further studies on probing potential functions of RBM20 in the mouse brain.

**Figure 1.**
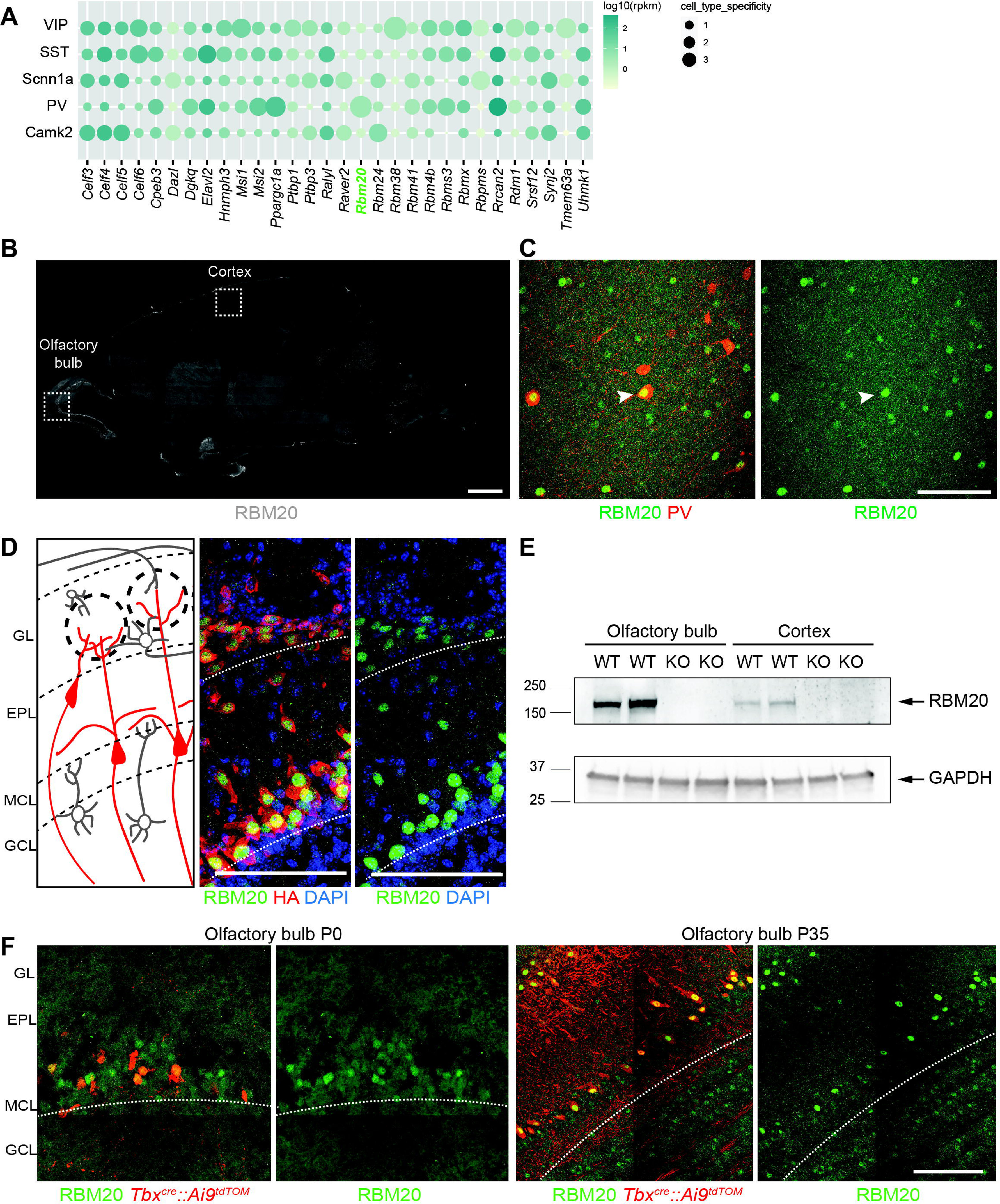
Characterization of *Rbm20* expression in the brain. A. Dot plot of the expression of a hand-curated list of RBPs across different neuronal neocortical populations. RBPs were chosen based on the presence of an RNA recognition motif (RRM) in their sequence and on the ranking of their gini-index value (only the first 20 RBPs displaying the highest gini-index value are displayed (see methods)). RBPs expression was measured by Ribo-TRAP sequencing and expressed as RPKM values normalized over the mean expression across different neuronal populations. B. Sagittal section of the mouse brain used for immunohistochemistry of RBM20 (grey). The somatosensory cortex and the olfactory bulb regions, where RBM20 is expressed, are highlighted. Scale bar: 1 mm. C. Immunohistochemistry of RBM20 expression (green) in Parvalbumin positive interneurons (red) in the neocortex. D. Schematic illustration of the olfactory bulb circuitry and cell types (left). GL: glomerular layer, EPL: external plexiform layer, MCL: mitral cell layer, GCL: granule cell layer, (left). RBM20 expression (green) is specific to the mitral cell layer and glomeruli layer of the olfactory bulb identified in the *Vglut2^Cre^ :: Rpl22^HA^*mouse line (HA staining in red) (middle and right). Scale bar 100 µm. E. Western blot probing RBM20 expression in olfactory bulb and cortex samples of wild-type (WT) and constitutive *Rbm20^KO^* mice (KO). The RBM20 band at ca. 150 kDa indicated with an arrow is selectively lost in KO tissue. GAPDH detection is used as loading control. For better visualization of the two proteins, the same tissue lysates were run on a 7.5% acrylamide gel (for RBM20 detection), and 4-20% acrylamide gradient gel (for GAPDH detection). F. Immunofluorescence of RBM20 (green) expression in the mitral cell layer (MCL) of P0 and P35 *Tbx21^Cre^::Ai9^tdTomato^*mice. Mitral cells and tufted neurons are labelled with the tdTomato reporter (red). Scale bar 100 µm.

To validate the neuronal expression, we investigated *Rbm20* mRNA distribution in the mouse neocortex by RNA fluorescent *in situ* hybridization (FiSH). This analysis confirmed significant expression of *Rbm20* in PV^+^ GABAergic interneurons (Fig. S1B-D). Moreover, we detected *Rbm20* mRNA in somatostatin-positive (SST^+^) interneurons of the somatosensory cortex (Fig. S1B - D). A survey of open-source datasets (Sjostedt *et al*, 2020) revealed that *Rbm20* mRNA expression in PV^+^ interneurons is evolutionary conserved in human. To examine RBM20 at the protein level, we raised polyclonal antibodies and confirmed RBM20 expression in PV^+^ interneurons of the somatosensory cortex (Fig. 1B and C). The parvalbumin-negative RBM20^+^ cells most likely represent somatostatin^+^- interneurons which also express significant levels of Rbm20 mRNA (Fig.S1C-D). Notably, these experiments also uncovered neuronal RBM20 expression outside of the neocortex, with particularly high protein levels in the olfactory bulb (Fig.1B,D, and E, see Fig.S1E for comparison to protein expression in the heart and Fig.S5A,B for antibody validation in immunostaining). In the olfactory bulb, RBM20 was restricted to glutamatergic cells in the mitral cell layer (MCL) and glomerular layer (GL) as opposed to the expression in GABAergic populations in the somatosensory cortex (Fig. 1D). Genetic marking and *in situ* hybridization for markers *Vglut2* and *Tbr2* confirmed the glutamatergic neuron expression of *Rbm20* mRNA in tufted and mitral cells (Fig. S2A-C), with only very low levels in a small number of GABAergic neurons (Fig. S2D). Similarly, we detected RBM20 protein in mitral cells of newborn mice (genetically marked with *Tbx^cre^::Ai9^tdTOM^*), suggesting that its expression is initiated during development and continues into the adult (Fig. 1F).

### Neuronal RBM20 binds distal intronic regions of mRNAs encoding synaptic proteins

In cardiomyocytes, RBM20 acts as an alternative splicing regulator that gates exon inclusion by binding to exon-proximal intronic splicing silencers (Li *et al*, 2013; Maatz *et al*., 2014; Dauksaite & Gotthardt, 2018). Within the nucleus, RBM20 localizes to foci in close proximity to *Titin* transcripts (Guo *et al*., 2012; Li *et al*., 2013) and has been proposed to form “splicing factories”, sites for coordinated processing of mRNAs derived from multiple genes (Bertero *et al*, 2019). By contrast, neuronal RBM20 did not concentrate in nuclear foci but instead, was distributed throughout the nucleoplasm (Fig. 2A). This difference in the sub-nuclear localization likely arises from the lack of *Titin* and/or other similarly abundant mRNA targets in neuronal cells. To contrast neuronal and cardiomyocyte RNA targets, we mapped transcripts directly bound by RBM20 in the olfactory bulb and in the heart using cross-linking immunoprecipitation followed by sequencing (CLIP-seq) analysis (Ule *et al*, 2003; Van Nostrand *et al*, 2017b). Our RBM20 antibodies lacked sufficient affinity for immunoprecipitation. Thus, we generated a *Rbm20^HA^* knock-in mouse line where the endogenous RBM20 protein is tagged with a Histidine-Biotin acceptor peptide and a triple HA-epitope (Fig. 2B and C and Fig. S3A and B). Introduction of this tag did not alter overall RBM20 protein levels or expression pattern (Fig. S3B, C).

**Figure 2.**
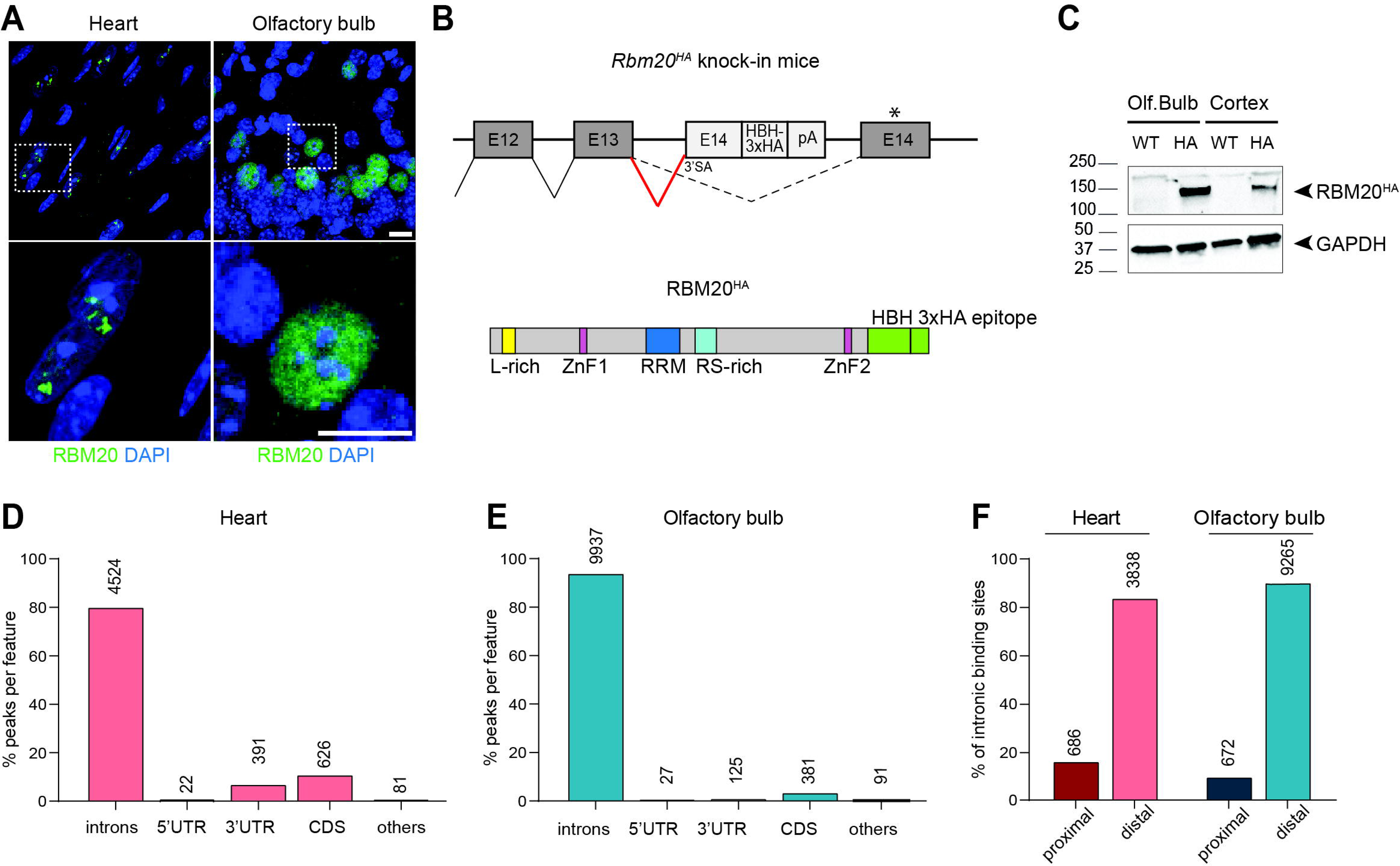
Identification of RBM20 direct targets in the heart and olfactory bulb. A. Sub-nuclear localization of RBM20 (green) in heart cardiomyocytes (left) and mitral cells of the olfactory bulb (right) of WT mice at P35. Scale bar 10 µm. B. Schematic illustration of HA epitope-tagging of endogenous RBM20 in mice. A cassette was inserted into the *Rbm20* locus containing a strong synthetic 3’ splicing acceptor site (3’SA) introduced into sequences derived from *Rbm20* exon 14 and in frame fusion of a histidine-biotin-histidine-3xHA tag, followed by a poly-adenylation signal (up). Schematic representation of the resulting RBM20^HA^ protein where the last exon of the protein is fused to a histidine-biotin-histidine-3xHA tag (down). C. Western blot showing the validation of RBM20 expression in the olfactory bulb and cortex tissues of WT and *Rbm20^HA^*tagged mice. GAPDH is used as a loading control. D, E (D) Quantification of the percentage of peaks identified in the heart (red) and (E) olfactory bulb (blue) tissue in each genomic feature: introns, untranslated regions (3’UTR, 5’UTR), coding sequence (CDS), others (promoters, intergenic regions, non-coding regions). The absolute number of the peaks identified in each genomic feature is reported on top of the corresponding bar. F. Bar plot showing the percentage of peaks identified in distal (>500 bp) or proximal (< 500 bp) intronic regions in the heart (red) and olfactory bulb (blue). The absolute number of the peaks identified is reported on top of the corresponding bar.

RBM20 CLIP-seq analysis on heart and olfactory bulb tissues from P35-P40 *Rbm20^HA^* knock-in mice identified significant peaks in 956 and 2707 unique transcripts, respectively. Recovery of protein-RNA adducts was UV-crosslinking dependent (Fig.S4A,B) and CLIP tags obtained in biological replicates were highly correlated (two replicates for heart and three for olfactory bulb, Fig. S4C and D-F). In both tissues, the majority of the identified RBM20 CLIP peaks mapped to introns (80% in the heart and 94% in the olfactory bulb) (Fig. 2D and E). Only a low fraction of peaks was identified in the 3’ untranslated region (11% in the heart and 1% in the olfactory bulb) and coding regions (7% in the heart and 3% in the olfactory bulb). RBM20 targets detected in the heart recapitulated all major target mRNAs identified in previous studies, including *Ttn, Tpm, Pdlim5, Scn5a, Camk2d, Cacna1c, Cacna1d* (Guo *et al*., 2012; Maatz *et al*., 2014; van den Hoogenhof *et al*., 2018; Watanabe *et al*., 2018; Fenix *et al*, 2021). In heart and olfactory bulb, approximately 90% of the intronic RBM20 binding sites localized to distal regions (> 500 bp from splice site) (Fig. 2F). The previously reported UCUU motif (Guo *et al*., 2012; Upadhyay & Mackereth, 2020) was significantly enriched at cross-link-induced truncation sites (CITS) (Fig. S4G) and *de novo* binding motif enrichment analysis identified a corresponding U-rich motif at RBM20 bound sites (Fig. S4H).

When directly comparing heart and olfactory bulb CLIP-seq datasets, we found transcripts from 363 genes that were commonly bound by RBM20 in both tissues. Other transcripts like *Ttn* are recovered as RBM20-bound only in one of the two tissues due to tissue-specific expression of the mRNA (Fig. 3A). Interestingly, *Camk2d* and *Cacna1c*, two important targets of RBM20 alternative splicing regulation in the heart, were identified as RBM20-bound in the olfactory bulb, however, binding sites mapped to different introns (Fig. 3A, and see TableS1). Finally, there was a sizable portion of pre-mRNAs (from 2721 genes) commonly expressed in both heart and olfactory bulb but selectively recovered as RBM20-bound in only one of the two tissues (539 genes in the heart and 2182 in the olfactory bulb). This suggests that RBM20 binds to target mRNAs in a tissue-specific manner. A tissue-specific function of RBM20 was further supported by gene ontology (GO) analysis. RBM20 target mRNAs in the heart showed enrichment for terms of muscle fiber components (M-band, Z-disc, T-tubule, Fig. 3B). By contrast, target genes in the olfactory bulb showed enrichment for terms related to pre- and postsynaptic structures, ion channels, and cytoskeletal components (Fig. 3C and Table S2). These include gephyrin (*Gphn*), a major scaffolding protein of GABAergic synapses, the adhesion molecule Kirrel3, and Downs Syndrome Cell Adhesion Molecule-like 1 (*Dscaml1*). This suggests that neuronal RBM20 plays a role in the regulation of synapse-related mRNA transcripts.

**Figure 3.**
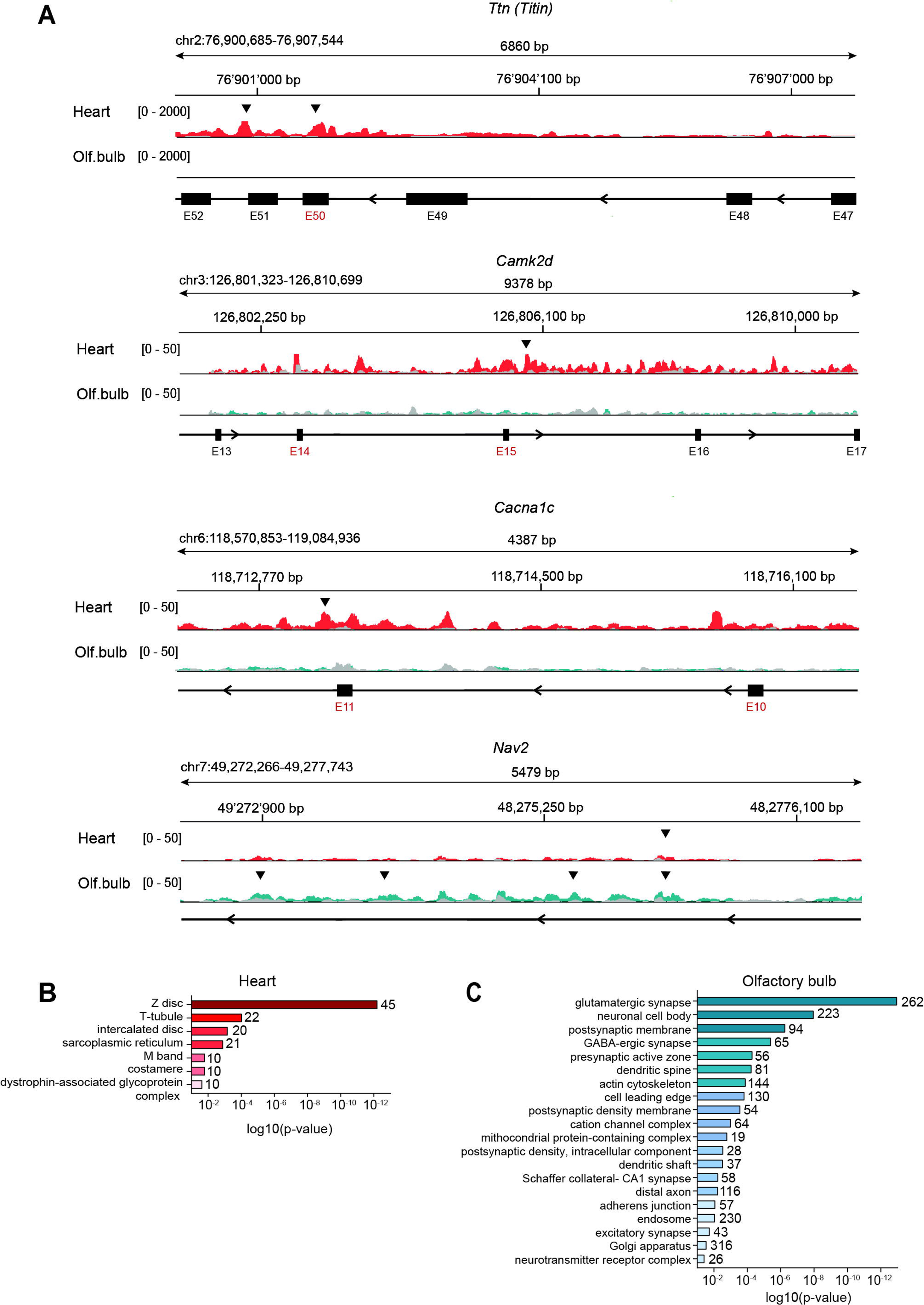
Identification of RBM20 direct mRNA targets in the heart and olfactory bulb. A. Tracks illustrating RBM20 CLIP-seq signal for *Ttn, CamkIId, Cacna1c,* and *Nav2*. Read density obtained for heart samples (red traces) and olfactory bulb (green traces) with the corresponding input samples (overlaid traces in grey). CLIP peaks, considered statistically significant by IDR (score >540, equivalent to IDR<0.05) are marked by black arrowheads. RBM20-dependent alternative exons previously reported for cardiomyocytes are labeled in red. Note that in the olfactory bulb, RBM20 binding sites are identified on *CamkIId* and *Cacna1c* pre-mRNAs. However, these binding sites are distal (> 500 bp) to the alternative exons. See TableS1 for coordinates. B. Illustration of the GO categories of RBM20 mRNA targets in the heart (IDR<0.05). Cellular Component analysis with Bonferroni correction (p-value ≤ 0.05). The number of genes found in each category is displayed on top of each bar. Minimal number of genes identified in each category: 5 genes. C. Illustration of the GO categories of RBM20 mRNA targets in the olfactory bulb (IDR<0.05). Cellular Component analysis with Bonferroni correction (p-value ≤ 0.05). The number of genes found in each category is displayed on top of each bar. Minimal number of genes identified in each category: 5 genes.

### Neuronal RBM20 is required for normal expression of long pre-mRNAs encoding synaptic proteins

To investigate the impact of RBM20 on the neuronal transcriptome, we performed loss-of-function experiments in *Rbm20* global and conditional knock-out mice. *Rbm20* was conditionally inactivated selectively in either parvalbumin-positive interneurons (*Pvalb^cre^::Rbm20^fl/fl^*, referred to as “*Rbm20^ΔPV^*”) or glutamatergic neurons (*Vglut2^cre^::Rbm20^fl/fl^*, referred to as “*Rbm20^ΔVglut2^*”). In global *Rbm20* knock-out mice, RBM20 immune-reactivity was abolished in the olfactory bulb and cortex (see Fig. 1E for western blots). Similarly, in the conditional knockout mice we observed a loss of RBM20 immune-reactivity in the respective cell populations (Fig. S5A and B). Importantly, neuronal cell types were normally specified in *Rbm20^ΔVglut2^* mice (Fig. S5C-F). We then applied a Translating Ribosome Affinity Purification protocol (RiboTRAP) optimized for small tissue samples (Heiman *et al*, 2014; Sanz *et al*, 2019; Di Bartolomei & Scheiffele, 2022) to uncover the impact of RBM20 loss-of-function on the neuronal transcriptome. Isolations were performed for *Rbm20^ΔPV^* and *Rbm20^ΔVglut2^* conditional knock-out and matching *Rbm20^WT^* mice (postnatal day 35-40). For quality control, we confirmed appropriate enrichment and de-enrichment of cell type-specific markers (Fig. S6A, B). Subsequently, we assessed the transcriptomes by deep RNA sequencing (4-5 replicates per condition, male and female mice, paired-end, 150=:Jbase pair reads, >80 million reads per replicate, see TableS3 and Fig. S6C - F for details). To identify genes with altered expression in *Rbm20* conditional knock-out mice we used DESEQ2 (Love *et al*, 2014). Using this approach, we did not identify significant differences in the overall transcriptome of *Rbm20^ΔPV^* PV^+^ interneurons as compared to wild-type (Fig. 4A and Table S4). However, in the olfactory bulb glutamatergic cells isolated from Rbm20*^ΔVglut2^* mice we identified de-regulation of 409 genes (FC ≥ 1.5, adjusted p-value < 0.01). 256 of these genes showed decreased expression in the conditional knockout mice as opposed to transcripts from 153 transcripts that were elevated (Fig. 4B). In the heart, RBM20 is considered a major regulator of alternative splicing. Thus, we quantified shifts in alternative exon usage upon *Rbm20* loss-of-function. We identified 859 differentially regulated alternative exons in PV^+^ interneurons and 1924 exons in the *Vglut2^+^* population in the olfactory bulb (FC in splicing index ≥ 1.5, p-value ≤ 0.05) (Fig. 4C,D and Table S5). The functions of the gene products with alternative splicing de-regulation were diverse and the only significantly enriched gene ontology term for genes with de-regulated exons was “mitochondrial protein-containing complexes” (p-value = 0.0003, 100 genes identified) for the olfactory bulb (*Vglut2^+^*) population (Table S6).

**Figure 4.**
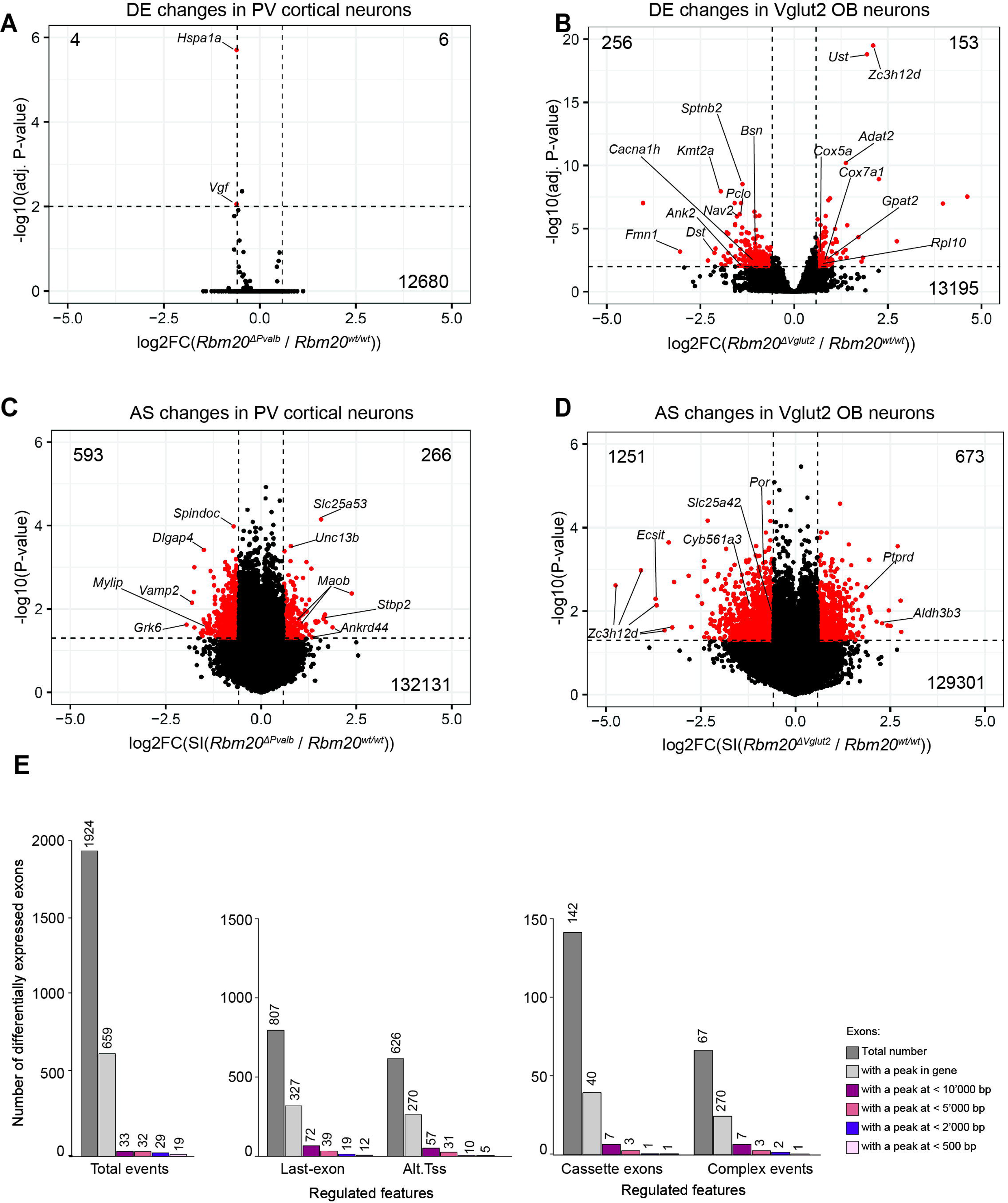
Differential gene expression and alternative exon incorporation rates in *Rbm20* conditional knock-out cells. A. Volcano plot of differential gene expression in RiboTrap-isolated mRNAs from *Rbm20*^WT^ *vs. Rbm20^ΔVglut2^* olfactory bulb (P35). Significantly regulated genes shown in red, cut-off fold-change (FC) of 1.5, adjusted p-value <0.01, total number of up- and down-regulated noted on top. Note that *Rbm20* itself is strongly reduced, outside the axis limits, and not represented in this plot (see Table S4). B. Volcano plot of the differential gene expression in RiboTrap-isolated mRNAs from *Rbm20^WT^ vs. Rbm20^ΔPV^* mouse neocortex (P35) as in panel A. The Y-chromosomal genes *Ddx3y*, *Uty, Kdm5d, Eif2s3y* were highly differentially expressed due to the larger number of *Rbm20* mutant males used in the Ribotrap isolations (3 wild-type females and 1 wild-type male vs. 4 knock-out male mice were used for this experiment). These genes and *Rbm20* itself were excluded from the plot (see Table S4 for complete data). C. Volcano plot representing differentially-included exons in *Rbm20^ΔVglut2^* RiboTrap-isolated mRNAs from olfactory bulb neurons. The dotted lines correspond to FC values of 1.5 and -1.5 and -log10 (p-value) of 1.3. Significantly regulated exons (FC 1.5 and p<0.05) are shown in red. D. Volcano plot representing differentially-included exons in *Rbm20^ΔPV^* RiboTrap-isolated mRNAs from cortical interneurons. The dotted lines correspond to FC values of 1.5 and -1.5 and -log10 (p-value) of 1.3. Significantly regulated exons (FC 1.5 and p-value <0.05) are shown in red. E. Number of exons differentially expressed in *Vglut2*^+^ cells isolated from the olfactory bulb of *Rbm20^ΔVglut2^* mice and number of exons with significant RBM20 CLIP peaks in binding sites within indicated distances. Equivalent information is provided for exons divided by annotations for specific features of regulation: alternative poly-adenylation (last exons), alternative transcription start sites (TSS), complex events and cassette exons.

To uncover transcript isoforms directly regulated by RBM20, we probed the intersection of the RBM20 binding sites identified by CLIP and alternative exon expression in the *Vglut2^+^* cell population. Of the 1924 exons with differential incorporation in *Rbm20^ΔVglut2^* cells, there were 659 exons with at least one significant RBM20 CLIP peak mapping to the corresponding gene. This frequency was higher than expected for a random distribution of CLIP peaks over all genes expressed in the sample (p = 5.2 * 10^-8^, hypergeometric distribution test for enrichment analysis). Interestingly, the vast majority of CLIP peaks were distant (>10kb) from differentially expressed exons. This applied regardless of the type of alternative exon feature (alternative last exon, alternative transcription start sites, cassette exons or complex alternative splicing events). Thus, we identified only 1 differentially regulated cassette exon with a proximal RBM20 binding event (< 500bp from the regulated exon) in the gene *Arhgef1*, (Fig. 4E). This suggests that suppression of alternative exons through proximal intronic splicing silencer elements is unlikely to be the primary essential function of neuronal RBM20.

Overall, RBM20 binding sites identified by CLIP distributed throughout the entire length of introns and did not exhibit a particular concentration at exon-intron boundaries (Fig. 5A). When examining the intersection of differential gene expression and CLIP data, we observed that 129 of the 256 down-regulated transcripts contained CLIP peaks (50.4%) whereas only 7 of the 153 up-regulated transcripts were bound by RBM20 (4.5%) (Fig. 5B). This suggests that RBM20 loss-of-function in neuronal cells results in the loss of mRNAs whereas the observed elevation of selective transcripts is likely to be an indirect, compensatory response. This notion is further supported by the distinct gene ontologies of the de-regulated transcripts. Down-regulated genes displayed a significant enrichment in GO terms related to synaptic components and the cytoskeleton (Fig. 5C and Table S6). By contrast, for the up-regulated genes, there was a pronounced enrichment in mitochondrial components and ribosome composition (Fig. 5C and see Table S6).

**Figure 5.**
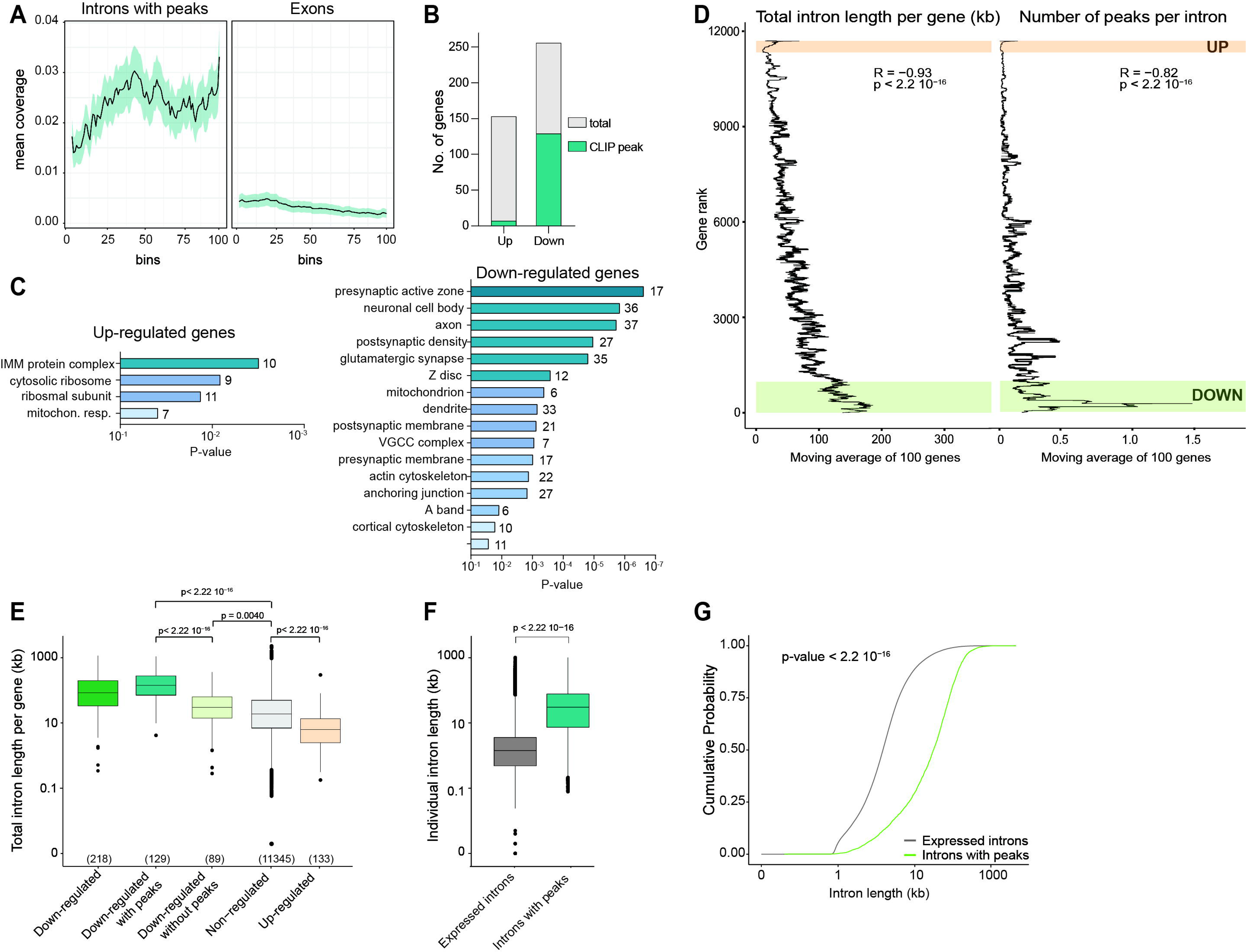
Long pre-mRNAs are depleted in *Rbm20^ΔVglut2^* mitral cells. A. Metagene coverage plots of CLIP peaks across all RBM20-bound introns. Peak density across exons from the same transcripts are shown for comparison. B. Total number of genes up- or down-regulated (FC > 1.5 and adjusted p-value <0.01) in glutamatergic cells from the olfactory bulb of *Rbm20^ΔVglut2^* mice. The fraction of genes with significant RBM20 CLIP peaks is indicated in blue. C. Illustration of gene ontologies enriched amongst up- and down-regulated genes (cellular component analysis with Bonferroni correction (p-value 0.05). Minimum number of 5 genes identified per category. D. Correlation of differential gene expression in *Rbm20^ΔVglut2^* cells, intron length and CLIP-seq data. Genes were ranked by FC in differential gene expression and mean total intron length (left) and mean number of intronic CLIP peaks (right) for blocks of 100 genes were plotted. Ranks of the genes meeting FC cut-off for down- or up-regulation are highlighted in color. Spearman’s coefficients and p-values are indicated. E. Boxplot showing the total intron length per gene (expressed in log_10_ scale) for categories of downregulated genes (all, with or without RBM20 binding sites), non-regulated genes, and up-regulated genes in *Rbm20^ΔVglut2^* in RiboTrap isolates from the olfactory bulb. Only annotated genes are plotted (number of genes shown at the bottom). P-values from Wilcoxon test are indicated. Medians: all down-regulated genes 83.2 kb, down-regulated genes with peaks: 152.4 kb; down-regulated genes without peaks: ∼33.6 kb; all non-regulated genes: 21.5 kb; all up-regulated genes: 7.4 kb. F. Boxplot (log_10_ scale) illustrating the total length of introns found in genes identified in our RBM20 Ribo-TRAP dataset (grey) compared to introns presenting RBM20 binding sites (green). RBM20 bound introns exhibit a higher intron length. P-values from Wilcoxon test are indicated. Medians: expressed introns: 1.4 kb; introns with peaks: 30.5 kb. G. Plot representing the cumulative probability distribution of intron length between the two groups of introns as in panel F. P-value (Kolmogorov-Smirnov test) is indicated.

TDP-43 and MATR3, two proteins with similar domain organization to RBM20, regulate target mRNAs by suppressing aberrant splicing into cryptic exons, a phenotype most significant for genes with long introns (Polymenidou *et al*, 2011; Ling *et al*, 2015; Attig *et al*, 2018). Interestingly, intron length of transcripts showed a significant correlation with differential gene expression and the number of CLIP peaks in *Rbm20^ΔVglut2^* cells (Fig. 5D - F). Importantly, mean intron length was substantially larger for down-regulated transcripts when compared to all detected genes or to up-regulated genes (Fig. 5E). This difference was amplified when selectively examining introns of down-regulated genes with CLIP peaks (Fig. 5F and G). Finally, introns bound by RBM20 were significantly longer than expected by chance as assessed with a permutation test. Random regions with the same properties as RBM20 CLIP peak regions were generated on introns from genes expressed in our Ribo-TRAP dataset (see methods). This resulted in a mean expected intron size of 41.5 kb which is substantially smaller than the mean size of RBM20-bound introns (59.0 kb; p-value < 0.0002). Thus, long neuronal mRNAs are preferentially bound by RBM20 and particularly sensitive to loss of neuronal RBM20 protein.

## Discussion

Our work establishes an unexpected function for RBM20 in neuronal RNA metabolism. We identified RBM20 expression in two specific neuronal populations in the mouse brain. Interestingly, these two populations are derived from highly divergent lineages: glutamatergic neurons of the olfactory bulb from the rostral telencephalon and PV^+^ GABAergic interneurons in the somatosensory cortex which arise from the medial ganglionic eminence of the subcortical telencephalon. Genetic deletion of RBM20 had only a modest impact on gene expression and transcript isoforms in PV^+^ interneurons but was associated with substantial alterations in glutamatergic cells of the olfactory bulb. This might be due to the higher expression of RBM20 in mitral and tufted cells as compared to PV^+^ interneurons. Moreover, PV^+^ interneurons express high levels of the RBM20 paralogue MATR3 (Fig.S1A) which may have overlapping functions.

In cardiomyocytes, several direct targets of RBM20-dependent alternative splicing regulation contain intronic RBM20 binding sites in close proximity to regulated exons (Maatz *et al*., 2014; van den Hoogenhof *et al*., 2018). In neuronal cells, we did not observe a similar regulation of proximal exons by RBM20. Interestingly, a large fraction of transcripts down-regulated in olfactory bulb RBM20 knock-out cells contained intronic RBM20 binding sites. Transcripts with long introns – which are prominently expressed in the mouse brain (Sibley *et al*, 2015; Zylka *et al*, 2015) – were particularly sensitive to RBM20 loss. Thus, in neuronal cells, the splicing repressive function of RBM20 might prevent the recruitment of cryptic splice acceptor sites within long intronic segments. We hypothesize that the reduced expression of RBM20 target mRNAs observed in *Rbm20^ΔVglut2^* neurons arises from aberrant nuclear splicing which ultimately result in the degradation of the transcripts.

The association of *RBM20* mutations with cardiomyopathies have directed substantial research efforts to uncover its function in the heart and to understand the impact of cardiomyopathy-associated mutations (Khan *et al*, 2016; Briganti *et al*., 2020; Schneider *et al*, 2020; Fenix *et al*., 2021). As for mice, human *RBM20* mRNA is also expressed in PV^+^ interneurons (Sjostedt *et al*., 2020). There is a growing body of literature highlighting shared genetic etiology of congenital heart disease and neurodevelopmental disorders (Homsy *et al*, 2015; Jin *et al*, 2017; Rosenthal *et al*, 2021). A significant fraction of children with congenital heart disease exhibit autistic traits including problems with theory of mind and cognitive flexibility (Marino *et al*, 2012) and the probability of an individual with congenital heart disease being diagnosed with autism was estimated to be more than twofold higher than in the typically developing population (Gu *et al*, 2023). A homozygous Ser529Arg substitution in the RNA recognition domain of RBM20 was recently identified in two brothers of consanguineous families affected by epilepsy and developmental delay (Badshah *et al*, 2022). The *RBM20* variant segregated with a second mutation in the *CNTNAP2* gene, which encodes a neuronal transmembrane protein implicated in axon-glia interactions. Thus, it remains to be explored whether alterations in neuronal RBM20 function contributes to the clinical characteristics observed in these individuals. However, given the neuronal expression and function of RBM20 identified in our study, a survey of – thus far unexplored – neurological phenotypes in *RBM20* mutation carriers might be warranted.

## Materials and Methods

### Immunochemistry and imaging

Animals (males and females) from postnatal day 25 to 40 were anesthetized with ketamine/xylazine (100/10 mg/kg *i.p.*) and transcardially perfused with fixative (4% paraformaldehyde). The brains or hearts were post-fixed overnight in the same fixative at 4°C and washed 3 times with 100 mM phosphate buffer. Coronal brain slices were cut at 40 µm with a vibratome (Leica Microsystems VT1000).

For immunohistochemistry, brain sections were processed as previously described (Traunmuller *et al*, 2023). In brief, brain slices were kept for 1.5 h with a PBS-based blocking solution containing 0.1% Triton X-100 and 5% normal donkey serum (NDS) and subsequently incubated with primary antibodies in blocking solution at 4°C overnight. Secondary antibodies were diluted in 5% NDS in PBS containing 0.05% Triton-X100 to a final concentration of 0.5 µg/ml or 1.0 µg/ml and incubated with sections for 2 hrs at RT.

Image stacks were acquired at room temperature on a laser-scanning confocal microscope (Zeiss LSM700) using a 40x Apochromat objective with numerical aperture 1.3, controlled by Zen 2010 software. Following acquisition, images were processed and assembled using Fiji (Schindelin *et al*, 2012), OMERO, and Adobe Illustrator software.

### Surgeries and stereotaxic injections

Recombinant Adeno-associated viruses with the rAAV2 capsid (Tervo *et al*, 2016) were produced in HEK293T cells. In brief, initially, a DNA mixture consisting of 70ug AAV helper plasmid, 70ug AAV vector, and 200ug pHGTI-adeno1 is prepared in a falcon tube and added at a 1:4 DNA:PEI ratio to each cell plate. After 48-60 hours, cells are collected and debris are collected by centrifugation at 4,000 rpm, 4°C for 20 minutes. The supernatant containing the virus is subsequently purified through OptiPrep^™^ Density Gradient Medium, (Sigma, Cat. No D1556). Viral preparations are concentrated in 100K Millipore Amicon columns at 4°C. The virus samples were then suspended in PBS 1X, aliquoted and stored at -80°C. Viral titers were determined by qPCR and were >10^12^ particles/mL.

Mice (postnatal day 24 to 27) were placed on a heating pad in a stereotaxic frame (Kopf Instrument) under isoflurane anesthesia. A small incision (0.5–1 cm) in the skin overlying the area of interest was made, and bilateral viral injections were performed in the posterior Piriform Cortex using a Picospritzer III pressure injection system (Parker) with borosilicate glass capillaries (length 100 mm, OD 1 mm, ID 0.25 mm, wall thickness 0.375 mm). Coordinates: ML = + 2,2 mm, AP = + 2,35 mm, DV = - 3,95 mm from Bregma. A volume of 100 nl of virus was delivered to each side, through repeated puffs over several minutes. Viruses used were rAAV2-CAG-DiO-eGFP or rAAV2-SYN-Cre virus, driving cre-dependent eGFP expression from the chicken beta actin promoter and human synapsin promoter, respectively. Ten days after viral infection, mice were anesthetized and transcardially perfused. Position of the viral injection site was confirmed on coronal brain slices using DAPI-stained sections on a AxioScan.Z1 Slide scanner (Zeiss) using a 20x objective.

### Tissue clearing and anatomical reconstruction of mitral cells

The olfactory bulb and part of the anterior prefrontal cortex of rAAV2-infected mice were dissected after transcardial perfusion and cleared using the Cubic L protocol (Tainaka *et al*, 2018). Olfactory bulbs were placed into a 5 ml Eppendorf tube filled with pre-warmed CUBIC L solution (10% N-butyl-di-ethanolamine and 10% Triton X-100 dissolved in MilliQ water). The tissue was incubated on a shaking plate at 37°C for 48 h. Cleared bulbs were washed in 50 mM PBS 3 times for 10 min and then cut into two coronal halves under a Binocular Stereo Microscope (Olympus #MVX10). The two halves of each bulb (anterior or posterior) were then embedded in 1% agarose in TBE solution in a glass bottom imaging chamber (Ibidi, Cat.No:80426). Z-stacks of GFP^+^ neurons with the soma residing in the mitral cell layer of the olfactory bulb were acquired on a Olympus two-photon microscope fitted with a MaiTai eHP laser (Spectra-Physics) and a 25X objective with 1.05 NA. A volume of up to 2 x 2 mm (xy) x 500-700 µm in depth was acquired in tiles up to 4x4, with x = 0.995 µm, y = 0.995 µm and z = 3 µm pixel size. Laser power was linearly adjusted with imaging depth and typically ranged between 0.5 to 20 mW. GFP^+^ mitral cells in the two-photon z-stacks were traced neurons using Neurolucida 360 software.

### Fluorescent *in situ* hybridization

Fluorescent *in situ* hybridization was performed using the RNAScope Fluorescent Multiplex Kit (Advanced Cell Diagnostics, Catalog Number 320851). P25 mouse brains were snap frozen in liquid nitrogen and 15 μm coronal sections were cut on a cryostat (Microm HM560, Thermo Scientific). Sections were fixed at 4°C overnight with 4% paraformaldehyde in 100 mM PBS, pH 7.4.

Images were acquired at room temperature with an upright LSM700 confocal microscope (Zeiss) using 40X Apochromat objectives (NA=1.3). Stacks of 10-15 µm thickness (0.44 µm interval between image planes) were acquired from layer 5 (L5) of the primary somatosensory area (S1). Genetically-marked cell types were identified based on the presence of transcripts encoding the *tdTomato* marker. Commercially available probes were used to detect *Rbm20* (ACD #549251) and *tdTomato* (ACD #317041). A region of interest (ROI) was drawn to define the area of the cell and dots in the ROI were manually counted throughout the image z-stacks. The number of dots in the ROI were then normalized to the cell area (measured in μm^2^). Images were assembled using Fiji and Adobe Illustrator Software.

### Biochemical procedures

Mouse tissues were extracted on ice and lysed in 50 mM Tris HCl pH 8.0, 150 mM NaCl, 0.1% SDS, 5mM EDTA, 1% Igepal, and protease inhibitor (Roche complete). The lysate was sonicated on ice (100 Hz Amplitude 0.5 cycles x 10 pulses) and centrifuged for 20 min at 13’000 g at 4°C. Proteins in supernatant were analyzed by gel electrophoresis on 4%–20% gradient polyacrylamide gels (BioRad, 4561093) and transferred onto nitrocellulose membrane (BioRad 1704158). Membranes were blocked with 5% non-fat dry milk (NFDM, PanReac AppliChem cat. no. A0830) blocking buffer in TBS-T 1X for 2h at RT and protein detection was by chemoluminescence with HRP-conjugated secondary antibodies (WesternBright Quantum, Advasta, Cat.no. K-12043 D20, K-12042 D20).

RiboTRAP purification (Heiman *et al*, 2008; Sanz *et al*, 2009) was performed with some modifications as described in (Di Bartolomei & Scheiffele, 2022).

### Library preparation and deep sequencing

For all the RNA-seq experiments, the quality of RNA integrity was analyzed using an RNA 6000 Pico Chip (Agilent, 5067-1513) on a Bioanalyzer instrument (Agilent Technologies) and only RNA with an integrity number higher than 7 was used for further analysis. RNA concentration was determined by Fluorometry using the QuantiFluor RNA System (Promega #E3310) and 50 ng of RNA was reverse transcribed for analysis of marker enrichment by quantitative PCR (*see* extended methods).

Up to five biological replicates per neuronal population and genotype were analyzed. Library preparation was performed, starting from 50ng total RNA, using the TruSeq Stranded mRNA Library Kit (Cat# 20020595, Illumina, San Diego, CA, USA) and the TruSeq RNA UD Indexes (Cat# 20022371, Illumina, San Diego, CA, USA). 15 cycles of PCR were performed. Libraries were quality-checked on the Fragment Analyzer (Advanced Analytical, Ames, IA, USA) using the Standard Sensitivity NGS Fragment Analysis Kit (Cat# DNF-473, Advanced Analytical).

The samples were pooled to equal molarity. The pool was quantified by Fluorometry using the QuantiFluor ONE dsDNA System (Cat# E4871, Promega, Madison, WI, USA) before sequencing. Libraries were sequenced Paired-End 151 bases (in addition: 8 bases for index 1 and 8 bases for index 2) setup using the NovaSeq 6000 instrument (Illumina) and the S1 Flow-Cell loaded at a final concentration in Flow-Lane loaded of 340pM and including 1% PhiX.

Primary data analysis was performed with the Illumina RTA version 3.4.4. On average per sample: 49±4 millions pass-filter reads were collected on this Flow-Cell.

### RNA Seq data analysis

Initial gene expression and alternative splicing analysis was done in collaboration with GenoSplice Technology, Paris as previously described (Furlanis *et al*., 2019). In brief, sequencing, data quality, reads repartition (e.g., for potential ribosomal contamination), and insert size estimation were performed using FastQC, Picard-Tools, Samtools and RSeQC tool packages. Reads were mapped using STAR (v2.4.0) (Dobin *et al*, 2013) on the mm10 Mouse genome assembly. The input read count matrix was the same as used for the splicing analysis. Two samples from the olfactory bulb were excluded from the analysis after the quality control of the data.

Read counts were summarized using featureCounts (Liao *et al*, 2014). For each gene present in the FASTDB v2021_4 annotations, reads aligning on constitutive regions (that are not prone to alternative splicing) were counted. Based on these read counts, normalization and differential gene expression were performed using DESeq2 (values were normalized to the total number of mapped reads of all samples) (Love *et al*., 2014) on R (v.3.5.3). Genes were considered as expressed if their FPKM value is greater than 96% the background FPKM value based on intergenic regions. A gene is considered as expressed in the comparison if it is expressed in at least 50% of samples in at least one of the 2 groups compared. Results were considered statistically significant for adjusted p-values ≤ 0.01 (Benjamini Hochberg for p-value adjustment as implemented in DESeq2) and log_2_(FC) ≥ 1.5 or or ≤-1.5. For the principal component analysis, counts were normalized using the variance stabilizing transform (VST) as implemented in DESeq2. The internal normalization factors of DESeq2 were used to normalize the counts for generation of heatmaps. The alternative splicing analysis was performed by calculating a splicing index (SI) which is the ratio of read density on the exon of interest and the read density on constitutive exons of the same gene. The log_2_ fold change (FC) and p-value (unpaired Student’s t-test) was calculated by pairwise comparisons of the respective Splicing Index (SI) values. Results were considered significantly different for p-values ≤ 0.01 and log_2_(FC) ≥1 or ≤-1.

For the generation of volcano plots, exons and genes with *NA* or *Inf* values were removed to prevent bias caused by genes or exons with very low expression. Plots were created in R with the *ggplot2* package.

### CLIP and library preparation

The CLIP experiments were performed according to the seCLIP protocol (Van Nostrand *et al*., 2017b) with some minor modifications (Traunmuller *et al*., 2023). Olfactory bulbs from seven mice were pooled for each biological replicate and for heart tissue one heart was used per biological replicate. Samples were flash frozen and ground on dry ice first in a metal grinder and a porcelain mortar. The frozen powder was transferred into a plastic Petri dish and distributed in a thin layer. The samples were UV-crosslinked 3 times at 400 mJ/cm^2^ on dry ice with a UV-crosslinker (Cleaver Scientific). The powder was mixed and redistributed on the Petri dish before each UV exposure. After crosslinking, the powder was collected in 3.5ml (olfactory bulbs) or 5.5 ml (heart) in lysis buffer (50 mM Tris-HCl pH 7.5, 100 mM NaCl, 1% NP-40, 0.1% SDS, 0.5% sodium deoxycholate, complete protease inhibitors, Roche) and 4 U per ml Turbo-DNase (Thermofisher). Samples were further processed as described in the extended methods section.

seCLIP data processing was performed as described (Van Nostrand *et al*, 2016; Van Nostrand *et al*, 2017a; Van Nostrand *et al*, 2020). In brief, raw reads were processed to obtain unique CLIP tags mapped to mm10 using Clipper (https://github.com/YeoLab/clipper; https://github.com/YeoLab/eclip). Reads from replicates 1 and 2 from the olfactory bulb were concatenated. Peak normalization was performed by processing the SMInput samples using the same peak calling pipeline. Irreproducible discovery rate (IDR) analysis was performed to identify reproducible peaks across biological replicates (Li *et al*, 2011). IDR (https://github.com/nboley/idr) was used ranking seCLIP peaks by the fold change over the size-matched input. Clip peaks were called based on IDR < 0.05. We observed some short highly represented sequences that were not specific to RBM20 seCLIP isolations, which were excluded based on peak shape and width (< 30 bp) using StoatyDive (Heyl & Backofen, 2021).

For motif discovery, crosslinking-induced truncation sites (CITS) were called using the CTK pipeline (Shah et al. 2017). Briefly, unique tags from replicates were combined and CITS were called by requiring FDR < 0.001. Sequences from -10bp to +10 bp from called CITS were used as input sequences for DREME software (Bailey *et al*, 2009; Bailey *et al*, 2015; Nystrom & McKay, 2021). As a control, sequences of the same length coming from 500 bp upstream of the (-510 to -490 bases) from the CITS site were used. Enrichment of the UCUU motif at the CITS sites was calculated.

### Analysis of RBM20-Bound intron length

For investigating the intron length of RBM20-bound introns, we performed a permutation test to calculate an empirical p-value. We generated 5,000 sets of random genomic coordinates, mirroring the length distribution and quantity of RBM20 seCLIP peaks. These coordinates were confined to the intronic regions of genes identified in the Vglut2 RiboTrap datasets.

In each permutation, we computed the mean length of all introns that included these random regions. The resulting distribution, based on 6,000 mean intron lengths, yielded a mean of ∼ 41.4 kb nucleotides and a standard deviation of ∼288.The average length of introns containing RBM20 seCLIP peaks was ∼ 59.0 kb, corresponding to a z-score of 60.95 and yielding a p-value 1.666667 * 10^-4^.This p-value is constrained by the number of permutations conducted.

### Gene Ontology analysis

All the gene ontology analysis were performed by using a statistical overrepresentation test and the cellular component function in PANTHER (http://pantherdb.org/). All genes being detected as expressed in the Ribo-TRAP RNA-sequencing data were used as reference. GO cellular component annotation data set was used and Fisher’s Exact test and Bonferroni correction for multiple testing was applied. GO terms with at least 5 genes and with P-value <0.05 were considered as significantly enriched. Significant GO terms were plotted in Prism 9.

For seCLIP GO analysis, any gene that had significant peak expression in the CLIP dataset either for olfactory bulb or heart samples was used and all genes being detected as expressed in the seCLIP size-matched input samples were used as reference.

### Statistical methods and data availability

Sample sizes were determined based on the 3R principle, past experience with the experiments and literature surveys. Pre-established exclusion criteria were defined to ensure success and reliability of the experiments: for stereotaxic injection, all mice with mis-targeted injections were excluded from analysis (e.g. if no eGFP signal was detected in the mitral cell layer of the OB). Investigators performing image analysis and quantification were blinded to the genotype and/or experimental group. For Ribo-TRAP pull-down experiments, all the samples presenting enrichment of the wrong marker genes were excluded. For the quantification of RBM20 expression in the olfactory bulb statistical analysis was performed with Prism 9 (GraphPad software) using unpaired t-test. Data presented are mean ± SD. Images were assembled using Fiji, Omero (Swedlow *et al*, 2003) and Adobe Illustrator software.

A detailed description of the exclusion criteria for different experiments is included in the respective method sections. Statistical analyses were conducted with GraphPad Prism 9. The applied statistical tests were chosen based on sample size, normality of data distribution and number of groups compared. Details on *n*-numbers, p-values and specific tests are found in the figure legends. All raw data files, excel analysis tables and additional data supporting the findings of this study could not be included in the manuscript due to space constraints but are available from the corresponding author upon reasonable request.

## Data availability

The datasets produced in this study are available in the following databases: MassIVE (code: MSV000093344), PRIDE (code: PXD046806), and GEO (code: GSE250100).

## Acknowledgements

We thank Caroline Bornmann and Sabrina Innocenti for excellent support with lab organization and experiments, Pawel Pelzcar and the Centre for Transgenic Models at the University of Basel for outstanding advice and services, Geoffrey Fucile (SciCORE) for help with data analysis, the Biozentrum Imaging Core Facility for support with image acquisition and analysis, the Quantitative Genomics Centre of the University of Basel for excellent technical assistance. The Scheiffele Laboratory is an associate member of the NCCR RNA & Disease, funded by the Swiss National Science Foundation. This work was financially supported by funds to P.S. from the Canton Basel-Stadt/University of Basel, the Swiss National Science Foundation (project 179432), a Swiss National Science Foundation Advanced Grant (TMAG-3-209273), and a European Research Council Advanced Grant (SPLICECODE).

