## Supplementary figures and images for "Dilated cardiomyopathy-associated RNA Binding Motif Protein 20 regulates long pre-mRNAs in neurons"

### Fig.S1

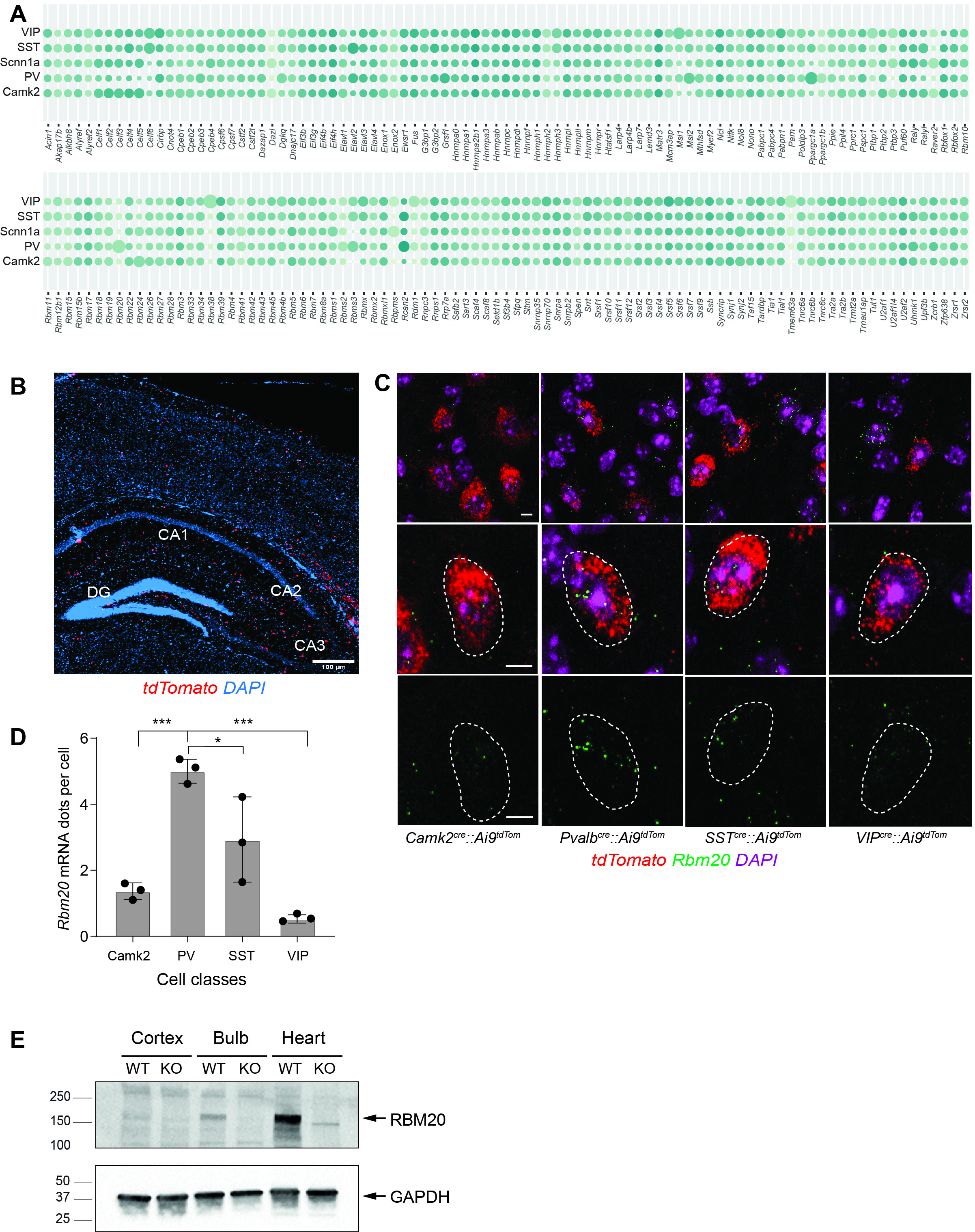

### Fig.S2

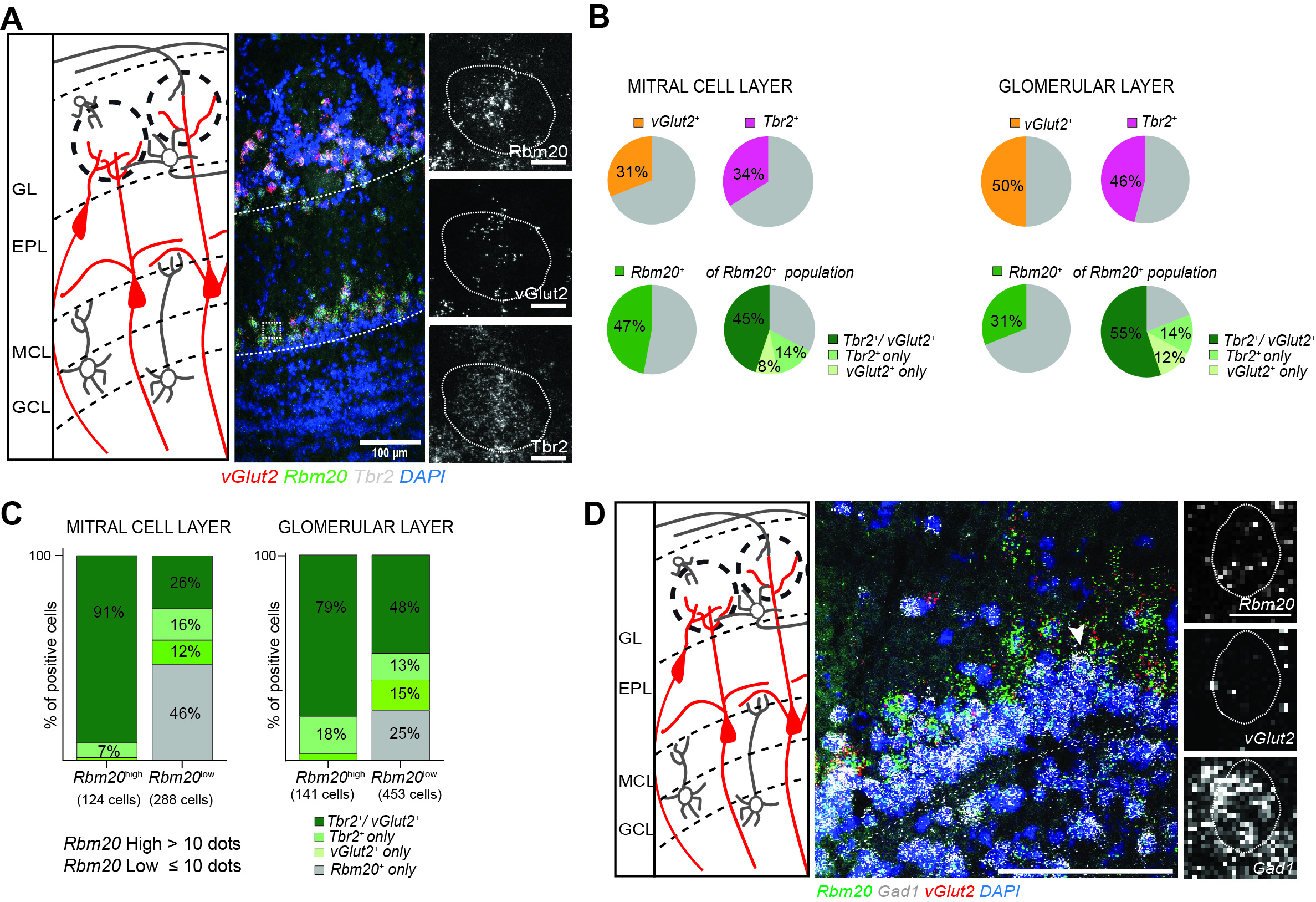

### Fig.S3

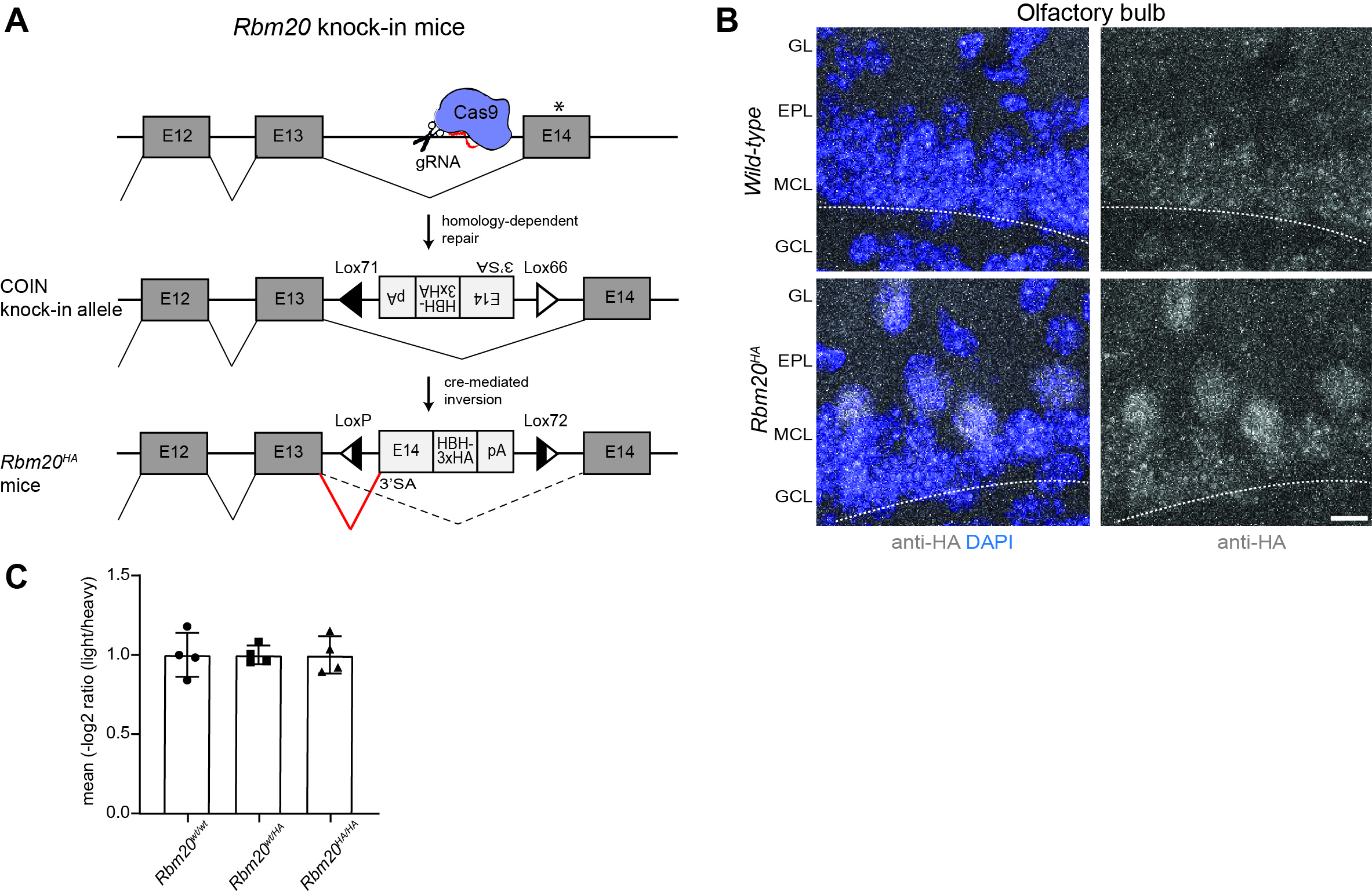

### Fig.S4

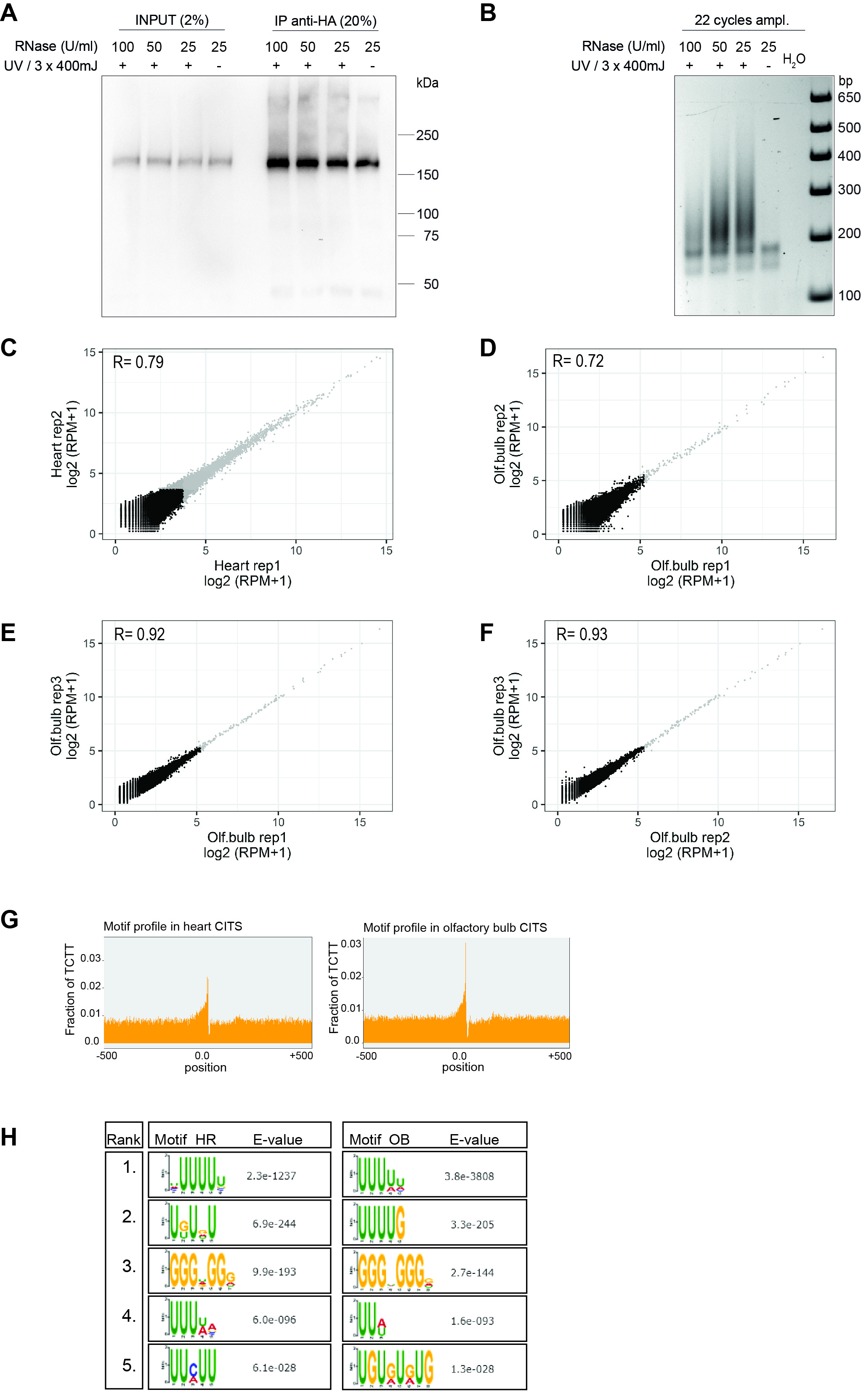

### Fig.S5

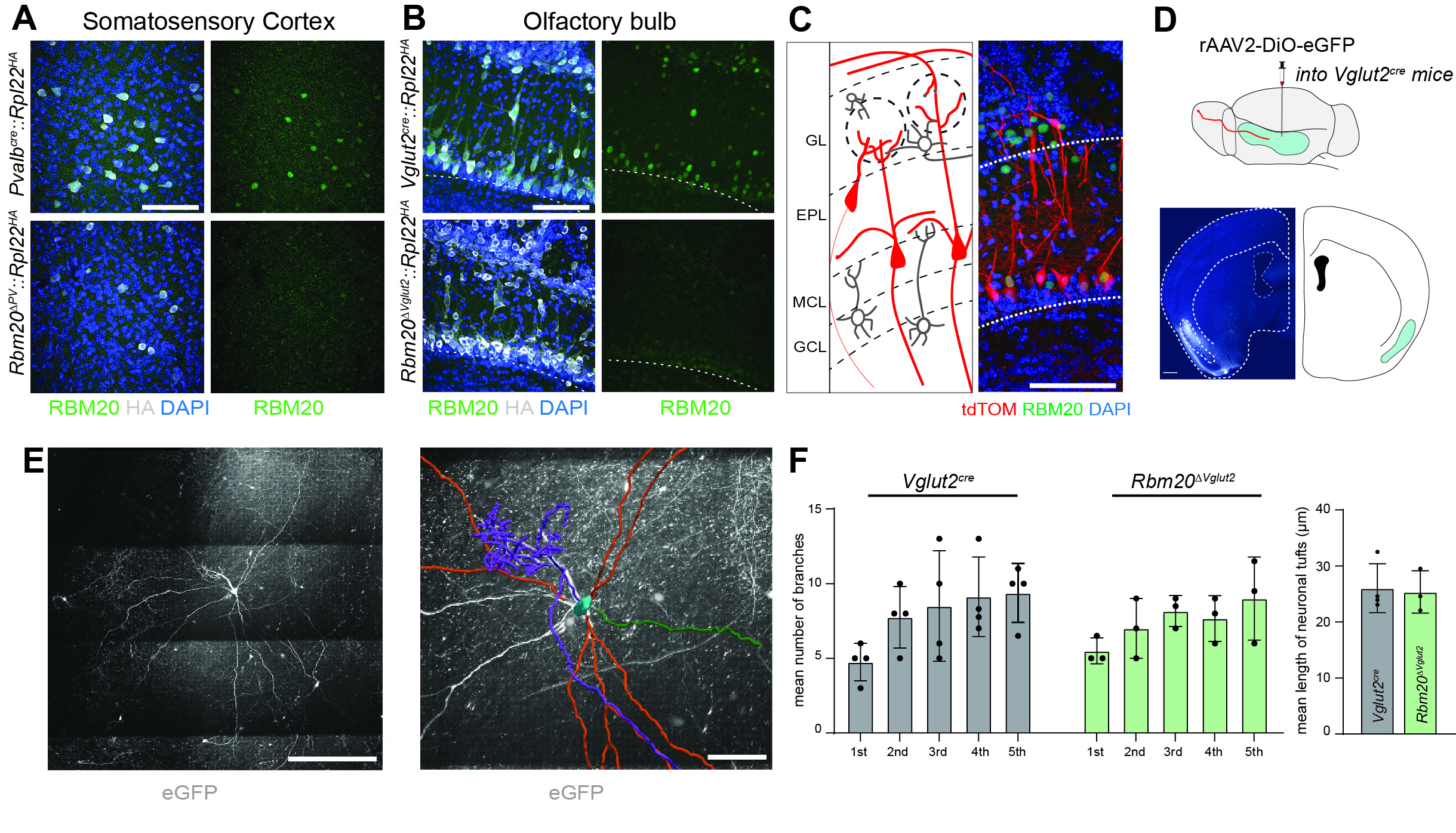

### Fig.S6

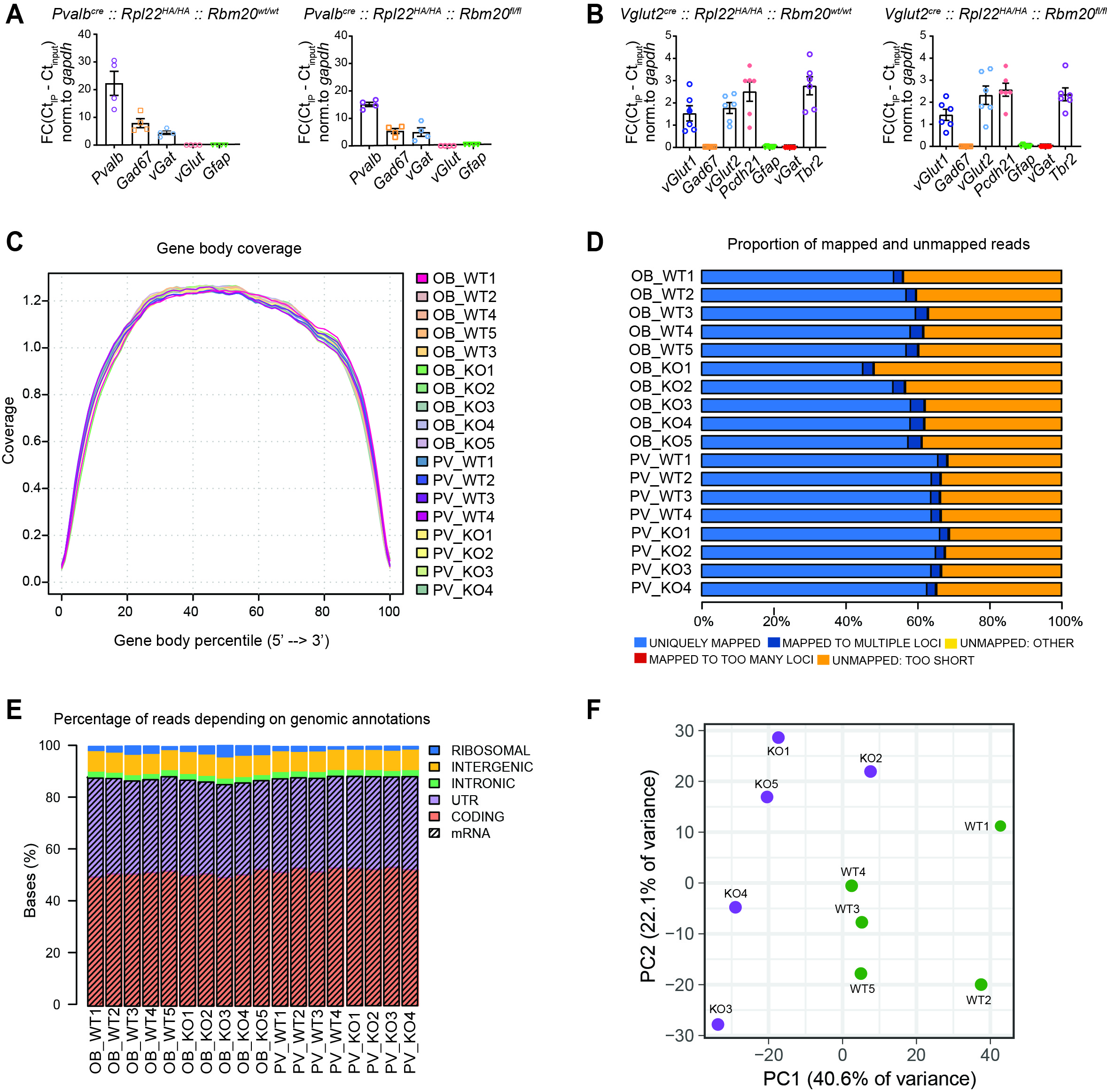
