## Supplementary text for "Dilated cardiomyopathy-associated RNA Binding Motif Protein 20 regulates long pre-mRNAs in neurons"

**Table of Content**

- Extended Materials and Methods
- Legend Figure S1 (related to Fig.1): Characterization of Rbm20 expression in the mouse neocortex
- Legend Figure S2 (related to Fig.1): RBM20 expression in mouse olfactory bulb
- Legend Figure S3, (related to Fig. 2 and 3): Generation of RBM20 – HA tagged mouse line
- Legend Figure S4, (related to Fig. 2 and 3): Identification of RBM20 binding sites on transcript mRNAs
- Legend Figure S5, (related to Fig. 4): Normal morphological differentiation of mitral cells in the absence of RBM20.
- Legend Figure S6, (related to Fig. 4): Quality control analysis of Ribo-TRAP RNA-sequencing samples
- Legend Table S1 (related to Fig. 2 and 3): Identified RBM20 binding sites in the heart and olfactory bulb tissues
- Legend Table S2 (related to Fig. 2 and 3): Gene Ontology analysis tables of transcripts directly bound by RBM20
- Legend Table S3 (related to Fig. 4): Number of expressed genes and percentage of mapped and unmapped unique reads in Ribo-TRAP RNA-sequencing experiments
- Legend TableS4 (related to Fig. 4 and 5): Summary tables resulting from differential gene expression analysis
- Legend TableS5 (related to Fig. 4 and 5): Summary tables resulting from the alternative exon usage analysis
- Legend Table S6 (related to Fig. 4 and 5): Gene Ontology analysis tables of de-regulated transcripts and alternatively spliced exons
- Legend Table S7 (related to Fig. 5): Intron length analysis table for de-regulated and non-regulated transcripts

**Extended Material and Methods**

**Mice**

All procedures involving animals were approved by and performed in accordance with the guidelines of the Kantonales Veterinärat Basel-Stadt. Male and female mice were used in this study. All other mice were in C57BL/6J background. *Rpl22^HA^* (RiboTag) mice (Sanz *et al*, 2009), *CamK2^cre^* (Tsien *et al*, 1996), *SST^cre^* (Taniguchi *et al*, 2011), *Pvalb^cre^* mice (Hippenmeyer *et al*, 2005), *Ai9^tdTOM^* reporter mice (Madisen *et al*, 2010), and *Vglut2^cre^* mice (Vong *et al*, 2011) were obtained from Jackson Laboratories (Jax stock no: 011029, 017320, and 007909, 028863, 007914 tdtomato and 016963). *Rbm20 ^floxed^* mice and *Rbm20* constitutive KO mice (Khan *et al*, 2016) were backcrossed for at least 4 generations to a C57BL/6J background. *Tbx21^cre^* mice (Haddad *et al*, 2013), were kindly provided by Dr. R. Datta’s Laboratory.

The *Rbm20* knock-in mouse line was generated in collaboration with the Center for Transgenic Mouse (CTM) lines in Basel. The line was generated using Crispr-Cas9. gRNAs targeting the last intron of *Rbm20* and template “*coin allele*” construct were injected together with RNA encoding for Crispr-Cas9 nuclease into C57BL/6J zygotes. The surviving embryos were transferred into recipient females. The *coin* module fused to a histidine-biotin-histidine-3HA tag. The *coin allele* is inserted in an orientation opposite to the gene’s direction of transcription. The gRNA used were: 5’ TTGAGTCGGGGGTCCCACTG 3’. The 1’311 bp single stranded megamer containing the upstream homology sequence (lower case), *coin module* (upper case letters) and downstream homology sequence (lower case) and the optimized codon usage (lower case) used:

5’ggcgaggctgctgctggagagccctgatttcttctctgtttgactcgcgaattctgaggggataagcgccctgcatatgtatgcattcttctttgggagcctgcagccaccttcatgcccagtaaggctatgcttactgtgccagatcaccccctgtaggctcacatagagccatgaccagcaacagcatagcgggatttccagaggcttcactgaggcagctatgacctgctcttgcctcccagggcatCCCCAGTACCGTTCGTATAatgtatgcTATACGAAGTTATGGGCCCCTCTGCTAACCATGTTCATGCCTTCTTCTTTTTCCTACAGAAGTACCTGTCTCAGCTGGCAGAGGAGGgactcAAGGAGACGGAGGGGACAGACAGCCCAAGCCCCGAGCGTGGTGGGATTGGTCCACACTTGGAAAGGAAGAAGCTAGCtGGcCAcCATCACCACCAcCATGGTGCcGCTGGAAAGGCCGGTGAAGGTGAAATCCCTGCCCCTCTTGCTGGTACaGTTTCTAAGATACTcGTAAAAGAAGGTGACACTGTTAAAGCTGGTCAAACAGTTCTGGTGCTGGAGGCcATGAAAATGGAGACAGAAATTAACGCTCCTACTGACGGAAAAGTTGAAAAGGTGTTAGTTAAGGAAAGAGATGCTGTTCAAGGTGGTCAAGGTCTAATCAAGATCGGCGTTGCAGGTCATCAcCACCAtCATCAcGGcGCcgccgggTATCCCTACGATGTGCCTGACTATGCTgctggcTATCCTTACGACGTGCCCGATTATGCAgccggcTATCCATACGATGTCCCAGATTACGCTgccTAGGATCTTTTTCCCTCTGCCAAAAATTATGGGGACATCATGAAGCCCCTTGAGCATCTGACTTCTGGCTAATAAAGGAAATTTATTTTCATTGCAATAGTGTGTTGGAATTTTTTGTGTCTCTCACTCGGAAGGACATATGGGAGGGCAAATCATTTAAAACATCAGAATGAGTATACCGTTCGTATAgcatacatTATACGAAGTTATTGGGACCCCCGACTCAAggtctcctgatgaatgctaactttctaagttgcctgacttgagtcagctggcacctgccctgtgggtcagacttcttcacttttcacacttgtggtttggagtaaagtgggagaggctgtagagactgaggcattcattctgccaaggcccctgacagaaacgctacctgagatggctgtggcagaggctcctggctccctgataaaaggtgtaccagggaaacgtgagctgaggtgggagggagtgagg’. The entire insert sequence is highlighted in capital letters. Activation by Cre-recombinase inverts the *coin* module, resulting in alternative splicing of the tagged exon. We observed only very low rates of cre-mediated inversion upon crossing to cre-driver mouse lines. Thus, a germline inverted allele was generated and used in all experiments. This constitutively tagged *Rbm20^HA^* mouse strain was deposited at EMMA.

All mouse lines were maintained on a C57Bl6/J background. Both males and females were used for all the experiments unless stated otherwise in the respective method sections.

**Antibodies**

The following commercially available antibodies were used: rat-anti-HA (Roche, #11867431001) and rabbit-anti-GAPDH (Cell Signaling #5174), rabbit-anti-MAP2 (Synaptic Systems #188002), rabbit-anti-RFP (Rockland #600-401-379), chicken-anti-GFP (Aves Labs Inc. #GFP-1020), goat anti-Parvalbumin (Swant, PVG213), rabbit-anti-NeuN (Novus Biologicals cat. n. NBP1-77686SS). Secondary antibodies coupled to horseradish peroxidase (HRP) or fluorescent dyes were from Jackson ImmunoResearch (goat anti-rabbit HRP #111-035-003; goat anti-rat HRP #112-035-143), donkey anti-rat IgG-Cy3 and Cy5 (Jackson ImmunoResearch, 712-165-153, 706-175-148) donkey anti-goat IgG-Cy3 and donkey anti-chicken IgG-Cy3 (Jackson ImmunoResearch, 705-165-147, 703-165-155).

For the generation of polyclonal anti-RBM20 antibodies, a synthetic peptide consisting of the RBM20 C-terminus was used: C+PERGGIGPHLERKKL (n- to c-terminus, C+ indicates a cysteine added to the n-terminus for thiol-mediated coupling). The synthetic peptide was conjugated to keyhole limpet hemocyanin for immunization of rabbits and guinea pigs (Eurogentec, Belgium). Resulting sera were affinity-purified on the peptide antigen and the specificity of the resulting antibodies was confirmed using lysates and tissue sections from *Rbm20* knock-out mice.

**Immunohistochemistry**

For quantifications of RBM20 positive neurons in the olfactory bulb, tile-scan images from 30 µm slices from the olfactory bulb of P35 mice were acquired. Mean intensity analyses for RBM20 signal were performed in Fiji (Schindelin *et al*, 2012) using a custom-made Python script, as previously described (DOI-https://github.com/imcf-shareables/3D_spots_count/blob/main/README.md). In brief, neuronal cells were identified based on the nuclear DAPI signal. The mean intensity of RBM20 protein in each nucleus was then measured and the background signal was subtracted.

For the characterization of RBM20 sub-nuclear localization, brain and heart samples from *Rbm20* WT and *Rbm20* cKO mice (P35-P40) were anesthetized with ketamine/xylazine (100/10 mg/kg *i.p.*) and transcardially perfused with fixative (4% paraformaldehyde). The brains and hearts were post-fixed overnight in the same fixative at 4°C and washed 3 times with 100 mM phosphate buffer (PB). Coronal brain slices were cut at 40 µm with a vibratome (Leica Microsystems VT1000). Brain samples were immersed in 15% and subsequently 30% sucrose in 1X PBS for 48 h, cryoprotected with Tissue-Tek optimum cutting temperature (OCT) and frozen at –80° until use. Tissue was sectioned at 40 µm on a cryostat (Microm HM560, Thermo Scientific) and collected in 1X PBS. Immunohistochemistry and imaging were performed as previously described.

**Targeted LC-MS sample preparation and analysis**

Murine nuclear extracts of heart tissue were lysed in 100 mM Triethylamonium bicarbonate pH 8.5 / 5% SDS / 10 mM Tris (2-carboxyethyl) phosphin using 20 cycles of sonication (30 s on / 30 s off per cycle) on a Bioruptor system (Dianode) followed by heating to 95° C for 10 min. Protein extracts were alkylated using 15 mM iodoacetamide at 25°C in the dark for 30 min. For each sample, 50 µg of protein lysate was captured, digested, and desalted using STRAP cartridges (Protifi, NY, US) following the manufacturer’s instructions. Samples were dried under vacuum and stored at –80°C until further use.

For parallel reaction-monitoring (PRM) assays (Peterson *et al*, 2012) three proteotypic peptides derived from RBM20 were selected for assay development (ASPPTESDLQSQACR, QGFGCSCR and SGSPGPLHSVSGYK). A mixture containing 100 fmol of each heavy reference peptide (JPT, Berlin, Germany) including iRT peptides (Biognosys, Schlieren, Switzerland) was used. The setup of the μRPLC-MS system was as described previously (Ahrne *et al*, 2016). Peptides were analyzed per LC-MS/MS run using a linear gradient ranging from 95% solvent A (0.15% formic acid, 2% acetonitrile) and 5% solvent B (98% acetonitrile, 2% water, 0.15% formic acid) to 45% solvent B over 60 min at a flow rate of 200 nl/min. Mass spectrometry analysis was performed on a Q-Exactive HF mass spectrometer equipped with a nanoelectrospray ion source (Thermo Fisher Scientific) as described previously (Hauser *et al*, 2022). The acquired raw-files were database searched against a *Mus musculus* database (Uniprot, download date: 2020/03/21, total of 44’786 entries) by the MaxQuant software (Version 1.0.13.13). To control for variation in sample amounts, the total ion chromatogram (only comprising peptide ions with two or more charges) of each sample was determined by Progenesis QI (version 2.0, Waters) and used for normalization. The datasets of this study are deposited on MassIVE (code: MSV000093344) and PRIDE (code: PXD046806).

**In situ hybridization**

For quantification of *Vglut2, Tbr2* and *Rbm20* transcripts expression in mitral and tufted neurons of the olfactory bulb, P25 animals were euthanized and the brains were harvested and processed as described above. Stacks of 10-15 µm width (0.44 µm interval between stacks) were acquired from olfactory bulb slices at room temperature with an upright LSM700 confocal microscope (Zeiss) using 40X Apochromat objectives. A ROI was drawn to define the area of each cell residing either in the mitral cell layer or glomeruli layer of the olfactory bulb. Dots in the ROIs were detected automatically throughout the z-stacks for each channel, using a custom-made Python script, as described in (DOI-https://github.com/imcf-shareables/3D_spots_count/blob/main/README.md). The following commercial probes were used: *Rbm20* (549251), *slc17a6* (319171), *Tbr2* (Eomes): (429641). Images from 3 mice were used for the quantification (2 images per slice). *Gad2* (415071) *in situ* hybridization was performed on 15 µm olfactory bulb slices of P25 mice.

**Two photon image analysis**

A total of 10 neurons (5 neurons per genotype from at least 3 biological replicates) were analyzed. Both apical and lateral dendrites of mitral cells were traced semi-automatically by using the user-guided 3D image detection algorithm. Tracings were checked and corrected manually when needed. Subcellular components, such as spines and other small protrusions were not traced. The following parameters were extracted from the software Neurolucida Explorer®. for each traced neuron: the number of dendrites from different centrifugal orders (*i.e.* primary dendrites, secondary dendrites etc.) and the total dendritic lengt of the glomeruli tufts. Average values were calculated for each neuron analyzed. Graphs and statistical analyses (t-tests) were made using GraphPad Prism.

**Quality control of ribotag pulldowns**

The enrichment and de-enrichment of markers following neuronal markers were tested: for the olfactory bulb pull-down samples: *Rbm20*, *Vglut2*, *Vglut1*, *Tbr2*, *Pcdh21*, Vgat, *Gfap*, *Gad67*. For pulldowns from cortex of PVCre mice: *Rbm20*, *Pvalb*, *Vgat*, *Gad67*, *Vglut1*, *Gfap*. In both cases, *Gapdh* mRNA was used as a housekeeping gene for normalization. The fold enrichment and de-enrichment values of each marker were calculated for each cell population in immunoprecipitated RNA, comparing it to input purifications. Only samples that showed correct enrichment or de-enrichment for excitatory or inhibitory neuronal markers and a de-enrichment for glia markers were further used for sequencing. DNA oligonucleotides were used with FastStart Universal SYBR Green Master (Roche, 4913914001) and comparative C_T_ method. For each assay, three technical replicates were performed and the mean was calculated. RT-qPCR assays were analyzed with the StepOne software. DNA Oligonucleotides used (name and sequence 5’-3’ are indicated):

List of primer sequences:

| Primer Name | Sequence |
| --- | --- |
| *Rbm20* | Forward (Sense)  TGCATGCCCAGAAATGCCTGCT  Reverse (AntiSense)  AAAGGCCCTCGTTGGAATGGCT |
| *Tbr2* | Forward (Sense)   ATAAACGGACTCAACCCCACC  Reverse (AntiSense)   CCCTGCATGTTATTGTCCGC |
| *Pchd21* | Forward (Sense)  ATCACTGTCAACGACTCAGACC  Reverse (AntiSense)  GTCAATGGCAGCTGAGTTTTCC |
| *Vglut2* | Forward (Sense)  GCATGGTCTGGTACATGTTCTG  Reverse (AntiSense)  GACGGGCATGGATGTGAAAAAC |
| *Gad67* \| | Forward (Sense)  GTACTTCCCAGAAGTGAAAC  Reverse (AntiSense)  GAATAGTGACTGTGTTCTAGG |
| *Gfap* \| | Forward (Sense)  CTCGTGTGGATTTGGAGAG  Reverse (AntiSense)  AGTTCTCGAACTTCCTCCT |
| *Vglut1* | Forward (Sense)  ACCCTGTTACGAAGTTTAACAC¨  Reverse (AntiSense)  CAGGTAGAAGGTCCAGCTG |
| *Vgat* | Forward (Sense)  CGTGACAAATGCCATTCAG  Reverse (AntiSense)  AAGATGATGAGGAACAACCC |
| *Pvalb* | Forward (Sense)  CATTGAGGAGGATGAGCTG  Reverse (AntiSense)  AGTGGAGAATTCTTCAACCC |

**CLIP sample preparation**

The lysate was transferred into a glass homogenizer and homogenized by 30 strokes on ice. 1 ml aliquots of homogenized tissue were transferred to 2 ml tubes, 10 µl of RNaseI (Thermofisher) diluted in PBS (1:5 -1:40) were added to each tube. Samples were incubated at 37°C with shaking for 5 min at 1200 x rpm and then put on ice. 10 µl RNasin RNase-inhibitor (40 U/µl, Promega) were added to each tube. Sample were mixed and centrifuged at 16.000 x g for 15min at 4°C. The supernatants were transferred to a new tube and 60 µl from each sample were taken and further processed for sized matched INPUT (SMIn). 10 µl HA-magnetic beads (Pierce) was added to each sample and incubated at 4°C for 4h in a rotating shaker. Following incubation, the beads were washed 2x with a high salt wash buffer (50mM Tris-HCl pH7.5, 1 M NaCl, 1 mM EDTA, 1% NP-40, 0.1% SDS, 0.5% sodium deoxycholate), 2x with the lysis buffer, 2x with low salt wash buffer (20 mM Tris-HCl pH7.5, 10mM MgCl_2_, 0.2% Tween-20) and 1x with PNK buffer (70 mM Tris-HCl pH6.5, 10 mM MgCl_2_). Beads were re-suspended in 100 µl PNK-mix (70 mM Tris-HCl pH6.5, 10 mM MgCl2, 1 mM DTT, 100 U RNasin, 1 U TurboDNase, 25 U Polynucleotide-Kinase (NEB)) and incubated at 37°C for for 20 min on a shaking termomixer (1200 x rpm). Upon RNA dephosphorylation, the beads were washed (2x high salt, 2x lysis and 2x low salt buffers as before) and additionally with 1x Ligase buffer (50 mM Tris-HCl pH7.5, 10 mM MgCl2). Beads were then re-suspended in 50 µl ligase mix (50 mM Tris-HCl pH7.5, 10 mM MgCl_2_, 1 mM ATP, 3% DMSO, 15% PEG8000, 30 U RNasin, 75 U T4 RNA-ligase (NEB)). 10 µl of the beads / ligase mix were transferred to a new tube and 1 µl of pCp-Biotin (Jena Bioscience) were added to validate IP of the RNA-protein-complexes by western blot. 4 µl of the RNA-adaptor mix containing 40 µM of each InvRiL19 & InvRand3Tr3 (IDT) were added to the remaining of the samples (40 µl). Samples were placed at RT for 2 h for adaptor ligation. Samples were washed 2x with high salt, 2x with lysis and 1x with low salt buffers. Finally, beads were re-suspended in 1x LDS sample buffer (Thermofisher) supplemented with 10 uM DTT and incubated for 10 min at 65°C, shaking on a thermomixer at 1200 x rpm. Eluates or inputs were loaded on 4-12% Bis-Tris, 1.5 mm gel (Thermofisher) and separated at 130 V for ~ 1.5 h. Proteins were transferred overnight at 30 V to a nitrocellulose membrane (Amersham). The membranes were placed in a 15 cm Petri dish on ice and an area between 55 and 145 kDa was cut out small pieces and transferred in a 2 ml tube.

RNA extraction, reverse transcription using InvAR17 primer, cDNA clean-up using silane beads (Thermofisher), second adaptor ligation (InvRand3Tr3) and cDNA purification steps were performed as previously described (Van Nostrand *et al*, 2016). The sequencing libraries were amplified using Q5-DNA polymerase (NEB) and i50X/i70X Illumina indexing primers (IDT). Final libraries were amplified with 14 cycles Libraries were purified and concentrated with ProNEX size selective purification system (Promega) using sample/beads ratio of 1/2.4. Samples were loaded on a 2% agarose gel and the area corresponding to the size between 175 bp and 350 bp was cut out. The amplified and purified libraries were then extracted from the gel using gel extraction kit (Machery&Nagel) and eluted with 16 µl.

The concentrations and the size distributions of the libraries were determined on the Fragment analyzer system (Agilent). 75 bp single-end sequencing was performed on the NextSeq500 platform using Mid Output Kit v2.5 (75 cycles).

Adaptor and primer sequences used in this study:

| Name | Sequence |
| --- | --- |
| InvRi L19 | /5Phos/rArGrArUrCrGrGrArArGrArGrCrArCrA rCrGrUrC/3SpC3/ |
| InvRand3Tr3 | /5Phos/NNNNNNNNNNAGATCGGAAGA GCGTCGTGT/3SpC3/ |
| InvA R17 | CAGACGTGTGCTCTTCCGA |
| i501 | AATGATACGGCGACCACCGAGATCTACACTATAGCCTACACTCTTTCCCTACACGACGCTCTTCCGATC*T |
| i502 | AATGATACGGCGACCACCGAGATCTACACATAGAGGCACACTCTTTCCCTACACGACGCTCTTCCGATC*T |
| i503 | AATGATACGGCGACCACCGAGATCTACACCCTATCCTACACTCTTTCCCTACACGACGCTCTTCCGATC*T |
| i504 | AATGATACGGCGACCACCGAGATCTACACGGCTCTGAACACTCTTTCCCTACACGACGCTCTTCCGATC*T |
| i701 | CAAGCAGAAGACGGCATACGAGATCGAGTAATGTGACTGGAGTTCAGACGTGTGCTCTTCCGATC*T |
| i702 | CAAGCAGAAGACGGCATACGAGATTCTCCGGAGTGACTGGAGTTCAGACGTGTGCTCTTCCGATC*T |
| i703 | CAAGCAGAAGACGGCATACGAGATAATGAGCGGTGACTGGAGTTCAGACGTGTGCTCTTCCGATC*T |
| i704 | CAAGCAGAAGACGGCATACGAGATGGAATCTCGTGACTGGAGTTCAGACGTGTGCTCTTCCGATC*T |

X* = Phosphorthioated base

**Supplementary Figures**

**Figure S1. RBM20 expression in mouse neocortex**

A.	Dot plot of the expression of a hand-curated list of RBPs across different neuronal neocortical populations. RBPs were chosen based on the presence of an RNA recognition motif (RRM) in their sequence and their expression in the neocortex. RBPs expression was measured by Ribo-TRAP sequencing and expressed in RPKM values normalized over the mean expression across different neuronal populations.

B.	Fluorescent *in situ* hybridization (FISH) on P25 brain slices from mice with genetic marking of cell populations (*Pvalb^cre^::Ai9* cre-dependent tdTomato expression). Red: *tdTtomato* mRNA, blue: DAPI. Scale bar 100 μm;

C.	Fluorescent *in situ* hybridization for *Rbm20* transcripts in *tdTomato* -marked cells for different cell classes (cre-dependent Ai9 tdTomato reporter crossed to the indicated cre-recombinase expressing lines: *Camk2*, *Pvalb*, *Sst*, *Vip*) in P23-26 somatosensory cortex cells in layer 5. Green: *Rbm20* mRNA, red: *tdTtomato* mRNA, blue: DAPI. Scale bar 10 μm.

D.	Quantification of *Rbm20* mRNA expression as in C, expressed as mean number of fluorescent dots per cell from three animals per genotype. P-value < 0.01, one-way Anova.

E.	Western blot analysis of RBM20 expression in cortex (CX), olfactory bulb (OB) and heart (HR) samples from WT and constitutive KO mice. A RBM20 antibody cross-reacting band with apparent mobility of ca. 150 kDa is lost in the KO samples (indicated with an arrow). Anti-GAPDH detection serves as a loading control.

**Figure S2. RBM20 expression in mouse olfactory bulb**

A.	Schematic illustration of the olfactory bulb circuitry and cell types. GL: glomerular layer, EPL: external plexiform layer, MCL: mitral cell layer, GCL: granule cell layer, (left). Fluorescent *in situ* hybridization (FiSH) on brain slices for *Rbm20* (green), *Tbr2* (gray), *Vglut2* (red) mRNAs (middle). The insets show the example of a cell expressing all three markers (right). Scale bar: 100 μm; scale bar insets: 10 μm.

B.	Pie charts indicating the quantification of the percentage of cells of the MCL and GL expressing *Vglut2, Rbm20* and *Tbr2* transcripts. Amongst the *Rbm20*^+^ cells, the percentage of neurons presenting co-localization with glutamatergic markers was calculated.

C.	Quantification of the percentage of neurons of the mitral cell layer (MCL) and glomerular layer (GL) expressing high or low *Rbm20* mRNA levels, co-localizing with *Tbr2* and *Vglut2* markers. The absolute number of high and low *Rbm20* expressing neurons identified in both the mitral cell layer and the glomeruli layer is reported in brackets.

D.	Fluorescent *in situ* hybridization (FISH) on brain slices of *Rbm20* (green), *Gad1* (gray), *Vglut2* (red) mRNAs. Scale bar 100 µm. The arrow in the inset on the right of the panel shows an example of a cell expressing low *Rbm20* levels and co-localizing with *Gad1* but not *V*g*lut2* mRNAs. Scale bar inset: 10 μm.

**Figure S3: Generation of *Rbm20* – HA tagged mouse line**

A.	Schematic illustration of the “COIN allele” strategy used for generation of *Rbm20^HA^* knock-in mice. The COIN allele module was inserted in the *Rbm20* locus with the CRISPR-CAS9 system. Upon Cre-mediated recombination of the loxP sites, the COIN module is inverted and the presence of a strong synthetic 3’ splicing acceptor site (3’SA) results in the expression of the tagged *Rbm20* isoform. We observed that the efficiency of cre-mediated inversion of this allele was too low for conditional tagging in cre-recombinase expressing mouse lines. Thus, an allele with germline recombination was generated resulting in constitutive tagging of RBM20 protein in all cells.

B.	Immunohistochemistry of RBM20^HA^ protein in mitral cells of wild type and *Rbm20^HA^* mice. HA (gray), DAPI (blue). Scale bar: 10 μm.

C.	Targeted proteomic analysis on heart samples of wild type (WT), *Rbm20^WT/HA^* heterozygous, and *Rbm20^HA/HA^* homozygous knock-in mice. Three proteotypic peptides were quantified. Plotted results represent means of all three peptides normalized to wild-type samples. Four biological replicates per genotype. The mean of the –log_2_ ratio (light/heavy) peptides was calculated and displayed. Note that an assessment of RBM20 expression level in the *Rbm20^HA^* mice by Western blot was not possible as the C-terminal HA-epitope reduced binding affinity of the anti-RBM20 antibody raised against the C-terminus of the protein.

**Figure S4: Identification of RBM20 binding sites on transcript mRNAs**

A.	Anti-HA Western blot for input (left) and immuno-precipitated HA-tagged RBM20 recovered from *Rbm20^HA/HA^* knock-in mice with precipitations performed after treatment with multiple RNaseA concentrations (25-100 units/ml) with (+) or without (-) UV-crosslinking of RNA-protein complexes.

B.	Agarose gel of libraries resulting from processed anti-HA IP samples amplified with 22 PCR cycles (using Illumina D701 and D501 primers). A negative control sample (water input instead of precipitated RNA) is shown on the right.

C-F.	Correlation analysis of normalized counts (reads per million [rpm] +1) of called CLIP peaks between seCLIP replicates of the heart (2 samples, panel A) and the olfactory bulb (3 samples, panels B,C,D). Gray shades represent density of the data points. Pearson coefficients are indicated in above the corresponding plots.

G.	Enrichment of the TCTT motif at cross-link-induced truncation sites (CITS) in both heart and olfactory bulb tissues. The enrichment is calculated from the frequency of the TCTT motif starting at each position of the inferred cross-linked site, normalized by frequency of the same motif in adjacent flanking regions.

H.	Motif finding analysis performed with DREME on heart and olfactory bulb seCLIP datasets. The first 5 statistically significant motifs are reported, ranked based on the enrichment p-value (E-value; right of each panel). The E-value is defined as the p-value times the number of candidate motifs tested. The enrichment p-value is calculated using Fisher's Exact Test for enrichment of the motif in the positive sequences, calculated after erasing sites that match previously found motifs. Note that the G-rich motif (rank 3) is commonly found to be non-specifically recovered in seCLIP datasets.

**Figure S5. Normal morphological differentiation of mitral cells in the absence of RBM20.**

A, B.	Immunostaining of RBM20 (green), RPL22-HA (gray) and DAPI (blue) in the cortex and olfactory bulb of P35 *Pvalb^Cre^::Rpl22^HA/HA^* and (B) *Vglut2^Cre^::Rpl22 ^HA/HA^* mice, compared to corresponding littermates carrying the conditional *Rbm20^fl/fl^* alleles (*Rbm20^ΔPV^* and *Rbm20^ΔVglut2^*, lower panels).

C.	RBM20-positive (green) mitral cells retrogradely labeled through injection of rAAV2-SYN-Cre virus (red) in piriform cortex of a *Ai9^tdTOM^* mice at P35. Scale bar 100 µm.

D.	Schematic illustration of the rAAV2-DIO-eGFP virus injection into the Piriform cortex of Vg*lut2^Cre^* mice for retrograde labelling of olfactory bulb mitral cells. Representative image of the site of viral injection in the piriform cortex. Scale bar 500 µm.

E.	GFP^+^ mitral cell (gray) retrogradely labeled through injection of rAAV2-SYN-DiO-GFP virus in the piriform cortex of Vg*lut2^Cre^* and *Vglut2cre::Rbm20^fl/fl^* (*Rbm20^ΔVglut2^*) mice. Tracing of the neuronal arborization is displayed on the right. Scale bar 500 µm.

F.	Quantification of the mean number of branches calculated per animal and the mean length of the neuronal tufts in reconstructed mitral cells of Vg*lut2^Cre^* and *Rbm20^ΔVglut2^* mice.

**Figure S6: Quality control analysis of Ribo-TRAP RNA-sequencing samples**

A – B.	Fold-enrichment (FC) of markers specific to inhibitory cortical neurons or glutamatergic neurons for WT and cKO samples. For PV samples, the following markers were tested: *Pvalb*, *Gad67*, *Vgat*, *Vglut1*, *Gfap*. For glutamatergic: *Vglut1*, *Vglut2*, *Gad67*, *Pcdh21*, *Gfap*, *Vgat* and *Tbr2*.

C.	Coverage plot indicating the percentage of read bases at a given position of the transcript. No sample displayed 3’ or 5’ coverage bias across the transcript length.

D.	Bar plot of each biological replicate where for each sample the following parameters are indicated: the proportion of number of reads uniquely mapped, mapped to multiple loci, mapped to too many loci or unmapped reads for all the samples. All samples show highly similar values across biological replicates, as well as across brain region and genotype, suggesting a high consistency and homogeneity of the RNA-seq data.

E.	Bar plot representing the relative percentage of reads falling on genomic features for all the biological samples.

F.	PCA of genes expressed in each olfactory bulb sample (n=5 biologically independent samples per genotype). Variance explained by the principal components 1 and 2 (PC1 and PC2) is indicated. Gene expression values were normalized by Variance Stabilizing Transformation (VST).

**Supplementary Tables:**

**Table S1: Identified RBM20 binding sites in the heart and olfactory bulb tissues.** List of RBM20 peaks identified on transcript mRNAs in the heart and in olfactory bulb tissues. Peaks were identified through the peak caller Clipper followed by irreproducible discovery rate (IDR) analysis between replicates. In this table, beyond the standard output parameters produced by IDR, columns containing information about the annotation of the targeted transcript and the position of RBM20 binding site in relation to the exon-intron boundaries (R package ‘AnnotatR) are reported. Moreover, the list of read counts summarized using featureCounts for identification of expressed genes in the input samples of heart and olfactory bulb is reported.

**Table S2: Gene Ontology analysis of transcripts directly bound by RBM20.** Gene ontology analysis results by Panther of mRNA transcripts directly bound by RBM20 RNA-binding protein in both heart and olfactory bulb tissues.

**Table S3: Expressed genes and percentage of mapped and unmapped unique reads in Ribo-TRAP RNA-sequencing experiments.** Read counts summarized using featureCounts. For each sample from either *Vglut2^+^* neurons of the olfactory bulb or PV^+^ interneurons of the neocortex.

**Table S4: Summary of differential gene expression analysis.** Expression values (FPKM) of genes identified in Parvalbumin positive and *Vglut2^+^* neurons in Ribo-TRAP RNA sequencing experiments and the results of their differential gene expression analysis by DESEQ2 in wild-type vs. *Rbm20* conditional knock-out mice.

**Table S5: Summary of alternative exon usage analysis.** Expression values (RPKM) of all the exons identified in Parvalbumin positive and Vglut2 positive cells in Ribo-TRAP RNA sequencing experiments and the results of the alternative exon usage analysis in wild-type vs. *Rbm20* conditional knock-out mice.

**Table S6: Gene Ontology of de-regulated transcripts and alternatively spliced exons.** Gene ontology analysis results by Panther for de-regulated transcripts (all, downregulated and upregulated) and exon usage analysis in olfactory bulb neurons upon RBM20 ablation.

**Table S7: Analysis of intron length.** Analysis of intron length in de-regulated, non-regulated and RBM20-bound transcripts in olfactory bulb neurons.
